# Neuroticism and emotion regulation: An effective connectivity analysis of large-scale resting-state brain networks

**DOI:** 10.1101/2023.04.21.537808

**Authors:** Tajwar Sultana, Muhammad Abul Hasan, Adeel Razi

**Affiliations:** Department of Computer and Information Systems Engineering, NED University of Engineering & Technology, Karachi 75270, Pakistan; Department of Biomedical Engineering, NED University of Engineering & Technology, Karachi 75270, Pakistan; Neurocomputation Laboratory, National Centre of Artificial Intelligence, Pakistan; Turner Institute for Brain and Mental Health, School of Psychological Sciences, Monash University, Clayton 3800, Australia; Wellcome Centre for Human Neuroimaging, University College London, WC1N 3AR London, United Kingdom; CIFAR Azrieli Global Scholars Program, CIFAR, Toronto, Canada

## Abstract

The neurotic personality has an impact on the regulation of basic negative emotions such as anger, fear, and sadness. There has been extensive research in search of functional connectivity biomarkers of neuroticism and basic negative emotions but there is a lack of research based on effective connectivity. In the current research, we intended to determine the significance of causal interaction of three large-scale resting-state networks – default mode, salience, and executive networks – to predict neuroticism and basic negative emotions. In this study, a large-scale human connectome project dataset comprising functional MRI scans and self-reported scores of neuroticism and negative emotions of 1079 subjects, was utilized. Spectral dynamic causal modelling and parametric empirical Bayes was used to estimate the subject-level effective connectivity parameters and their group-level associations with the neuroticism and emotional scores. Leave-one-out cross-validation using parametric empirical Bayes was employed for prediction analysis. Our results for heightened emotions showed that the self-connection of right hippocampus can predict individuals with high fear, self-connections of dorsal anterior cingulate cortex, posterior cingulate cortex and left dorsolateral prefrontal cortex can predict individuals with high sadness. High anger, low sadness, and neuroticism scores of any emotion category except low fear, could not be predicted using triple network effective connectivity. Our findings revealed that the causal (directed) connections of the resting-state triple network can potentially serve as a connectomic signature for people with high and low fear, high sadness, low anger, and neuroticism with low fear.

## Introduction

There is a growing interest in identifying the neural bases of personality traits because of the potential impact of personality on mental and physical health. A person’s personality is made up of consistent behavioral, cognitive, emotional, and motivational patterns that define them as a unique individual. The five-factor model or Big Five is currently the most extensively used taxonomy of human personality traits. It consists of five personality traits: openness, conscientiousness, extraversion, agreeableness, and neuroticism. Openness is about being creative and curious, conscientiousness is about being thoughtful, extraversion is about being social and active, agreeableness is about being cooperative and empathetic,, and neuroticism is about experiencing negative emotions or emotional instability [1], [2]. Among the five personality traits, neuroticism has the most relevance to a person’s mental and physical well-being [3]–[8]. Earlier research has shown that highly neurotic personalities are associated with dysregulation of negative emotions [9], [10]. Neurotic people are more prone to negative emotions such as anger, fear, anxiety, guilt, sadness, and depression. A study after controlling demographic factors revealed that neuroticism has a positive association with all the dimensions of mental health including social dysfunction and anhedonia, depression and, anxiety and loss of confidence while other four personality traits were associated with subsets of the mental health dimensions [7].

The importance of identifying and mapping the neural correlates of neuroticism is already established through many brain functional and effective connectivity research studies, as high neuroticism is a risk to many mental disorders including substance abuse and mood disorders [11]– [14]. Neuroticism is associated with executive [15], [16], memory [17], [18], affective [19], [20] and social functioning [21]. The relationship of neuroticism with other mental faculties entails the significance of resting-state networks involved in their processing to understand the neuroticism dynamics. The intrinsic dynamics of brain networks may better reflect these functions. This is the reason many researchers have employed resting-state brain connectivity techniques to unveil the mechanisms of neuroticism. There have been many resting-state studies to understand the intrinsic brain function associated with trait neuroticism. The investigation of the associations of neuroticism with resting-state fMRI data have yielded negative association of ventral and dorsal attention networks [22], diminished dmPFC and amygdala FC during negative emotion regulation [10], increased effective connectivity from amygdala to middle frontal gyrus and decreased effective connectivity from precuneus to amygdala [23], association with grey matter clusters of dorsal anterior cingulate cortex and medial prefrontal cortex [24] and so on. In highly neurotic individuals, external motivational factors alters the brain connectivity, such as criticism alters the functional connectivity between brain regions that are involved in negative emotions expression and regulation [25] and negative picture viewing decreased the functional connectivity between amygdala and frontal brain regions in high neurotic subjects [26].

Neuroticism is a heterogeneous trait with multiple facets including anger, sadness, anxiety, worry, and hostility [27]. There are four basic human emotions - happiness, fear, anger and sadness [28]. Emotions are the subjective feelings or thoughts of an individual which usually arise from past experiences and impact their behavior influencing general health and well-being. Negative emotions are shown to be positively associated with psychological health [29] and their improper regulation leads to mental health disorders [30], [31] and physical illnesses [32]. Neuroticism is shown to have a positive correlation with the duration and frequency of the negative emotions [33]. Effective regulation of negative emotions is essential for psychological well-being and this capacity is disrupted in highly neurotic individuals. The response to the aversive environmental conditions or surrounding situation arises due to negative emotions of anger, fear, and sadness. All other negative emotions are secondary or a result of these three primary or basic negative emotions – anger, fear, and sadness. Therefore, to fully comprehend the neural mechanisms of neuroticism, it is necessary to understand the neural mechanisms of fine-grained neurotic aspects i.e., the basic negative emotions. There are many brain functional and effective connectivity studies of basic negative trait affects. Functional connectivity association has been explored with the self-reported negative trait emotions [34]–[39]. Recently, a study was conducted to examine the effective connectivity associated with trait somatic anxiety which yielded that there was improved top-down effective connectivity at high fear-related somatic arousal [40]. Researchers have also predicted negative trait affects using functional connectivity [35], [41]. Previously, the brain connectivity of each of the basic negative emotions has been extensively studied but not together. However, there is a need to investigate their effective connectivity collectively as intrinsic processes during the resting-state i.e., without any external stimulation.

The most popular instruments for measuring neuroticism (and negative emotions) are self-reported questionnaires that involves questions representing continuous dimension and are generalized such as “Do you get irritated quickly” [42]. Self-reports are also a popular measure to assess both positive and negative emotions of an individual. One such example is NIH toolbox emotion battery that is a standard assessment tool designed to measure the positive and negative emotions. Their scoring system depends on item response theory (IRT) and IRT is based on the assumption that individuals will possess some extent of the underlying trait and will respond in a peculiar way [43], [44]. Therefore, the scores of NIH toolbox emotion battery discloses trait-like personality of the individuals related to emotions.

Resting-state functional connectomic fingerprints have been shown to identify an individual in a group though their functional connectivity [45]. Many researchers have also employed different techniques to predict a neurotic individual using their resting-state functional connectivity such as Nostro et al., showed that resting-state functional connectivity of 9 resting-state networks spanning different mental faculties including social, cognition and affection could predict neuroticism [46]; another study showed the utilization of resting-state functional connectivity to predict neuroticism through 244 brain regions organized into 8 canonical networks, through a machine learning algorithm – connectome-based predictive modelling (CPM) [47]. Recently, a personality prediction study based on resting-state functional connectivity of HCP subjects revealed that neuroticism can be predicted using functional connectivity of DMN, frontoparietal network, somatosensory network, visual network and cerebellum through CPM [48], later they inferred that personality traits including neuroticism are associated with resting-state functional connectivity of DMN and autonomic nervous system but CPM could not predict neuroticism significantly using these networks functional connectivity [49]. Another recent study showed that neuroticism can be predicted using resting-state functional connectivity of DMN and dorsal attention/sensory network as they showed less variation in functional connectivity profiles over a timespan [50]. On the contrary, another functional connectivity study based on graph theory that also used HCP dataset, declared no association of 15 resting-state networks including DMN, SN and frontoparietal network, with neuroticism [51]. Liu et. al., also used HCP data and found that dizygotic or non-identical twins have similar personality including neuroticism and similar personality-related functional connectivity patterns across many regions of resting-state networks including less studied visual network and cerebellum [52]. They also predicted varied facets of personality of an individual using functional connectivity. Although there are various functional connectivity-based prediction studies on neuroticism, functional connectivity is based on correlation analysis and hence cannot establish the directed connectivity between brain regions.

In this study, we are to investigate the association of neuroticism and negative emotions with the brain effective connectivity of three large-scale resting-state networks – default mode network (DMN), executive network (EN) and salience network (SN). Our objective is to determine the significance of the causal interaction of the triple network regions in predicting negative emotions and neuroticism. The salience network (SN) is the most pronounced resting-state network in the affective neuroscience. The reason being that SN comprises of limbic regions (such as amygdala) and cortical regions (such as insula and cingulate cortex) that are the major hubs of emotion processing. Although SN is the most prominent in emotional processing, it does not work in isolation, rather it interacts with other brain networks to process emotions. The most popular trio among the resting-state networks is DMN, SN and EN, where SN modulates the activity of DMN and EN regions and works as a modulating switch between the two networks. DMN is the most widely studied resting-state network [53]. The brain regions comprising DMN exhibit correlated increased brain activity when the brain is not engaged in any external task or during day dreaming or mind wandering [54]. DMN has shown to be involved in various phenomenon such as self-reflection [55], social [56] and emotional processing [57]. The role of DMN in emotional processing is mainly due to its function in internally generated thoughts including self-reflection, imagination, past ruminations, planning and thinking about others perspectives [58]–[60]. Specifically, the roles of mPFC and PCC in self-reflection, and hippocampus in emotional memory processing are well-established. The executive network is mostly involved in attention-seeking or goal-oriented tasks. It majorly comprises of dorsolateral prefrontal cortex (DLPFC) which is involved in the cognitive processing and valence regulation of emotions [61].

This study utilizes a large-scale resting-state fMRI dataset of 1079 subjects along with their NIH battery of emotions scores that include a subdomain of self-report basic negative emotions and NEO Five factor inventory (NEO-FFI) scores that provides measures of five personality domains including neuroticism. Considering the role of each of these three networks in emotion processing, we hypothesize that the causal (directed) connections between their regions would be able to predict neuroticism and negative emotions. This may serve as the neural signature for subjects with heightened emotion and neuroticism and may guide stimulation targets for mental health therapies.

## Methods

### fMRI Dataset

The resting-state fMRI scans and emotional function data of 1079 healthy young adults (females = 584, age: 22-35 years, mean age: 28.78 ± 3.5) was derived from the Human Connectome Project (HCP) [62]. The resting-state fMRI data was collected using Siemens 3T scanner. The acquisition parameters included TR=720 ms, TE=33.1 ms, 1200 frames per run, flip angle = 52 deg, FOV = 208×180 mm. The data was already preprocessed using fMRI volume minimal preprocessing pipeline of HCP [63]. The head motion and physiological noise correction was performed using nuisance regression, with six parameter estimates (translational and rotational) from rigid body transformation (downloaded from HCP) and CSF and WM timeseries as regressors.

### Emotions measures

The emotion measures consist of NIH Toolbox Emotion Battery [43]. Only self-report emotional function measurements are included in the NIH Toolbox. It consists of positive and negative subdomains of emotional health including measures of negative affect (anger, fear, and sadness), psychological well-being (positive affect, general life satisfaction, meaning and purpose), stress and self-efficacy (perceived stress and self-efficacy) and social relationships (social support, companionship, social distress). Our focus was on negative emotions, therefore utilized only scores of negative affect subdomain.

NIH Toolbox measures anger using anger-affect survey, anger-hostility survey, and anger-physical aggression survey. The self-report measure of anger-affect assesses anger as an emotion while anger-hostility and anger-physical aggression assesses hostility and aggression. Likewise, the toolbox measures fear using fear-affect survey and fear-somatic arousal survey which assesses fear as an emotion and somatic symptoms respectively. We used only relevant scores of anger-affect and fear-affect. The participants were asked questions about their emotional experience during the past 7 days prior the scan. Each emotion measurement was assessed using a computerized adaptive test (CAT) comprised of 4-6 items or questions derived from a question bank. The participant responds to each item by selecting one out of five response choices with raw scores 1 to 5 where high score denotes a strong emotion. The raw scores are summed up and converted into standard T-scores that are based on US healthy population. For this scale, the mean T-score is 50 with a standard deviation of 10. The scores below the mean (T ≤ 40) indicate low levels while above the mean (T ≥ 60) indicate high levels of fearful, angry, and sad feelings.

### Personality measures

The personality measures consist of NEO five-factor inventory (NEO-FFI) [64] in the HCP behavioral data. It is a self-reported questionnaire comprising of 60 items, 12 for each of the five personality factors – neuroticism, openness, extraversion, conscientiousness, and agreeableness [2]. Participants report their level of agreement for each item, on a five-point Likert scale. The summary scores for each personality trait are obtained by summing the codes for each answer (strongly disagree = 0, disagree = 1, neither agree nor disagree = 2, agree = 3, strongly agree =4). Some items are reverse coded appropriately. We only used the total scores for neuroticism.

### Regions of interest selection

Regions of interest (ROIs) spanning the three large-scale resting-state networks, default mode network (DMN), salience network (SN) and executive network (EN), were selected based on their significance in the emotion processing and emotion regulation. The chosen ROIs are posterior cingulate cortex (PCC), medial prefrontal cortex (mPFC) and bilateral hippocampus (lHP and rHP) for the DMN, dorsal anterior cingulate cortex (dACC), bilateral anterior insula (lAI and rAI) and bilateral amygdala (lAMG and rAMG) for the SN, and bilateral dorsal lateral prefrontal cortex (lDLPFC and rDLPFC) for the EN. The regional centroids were identified using associativity test maps of the regions from neurosynth.org. The associativity test maps consist of the z-scores indicating the non-zero association between the ROI and voxel activation. These maps were utilized to locate the centroid of each ROI by identifying the local maxima of the selected cluster in the map (using xjview [https://www.alivelearn.net/xjview]). The spherical regions of 8 mm were selected with the respective centroids (Table 1). To avoid the spheres, of smaller-sized regions, from incorporating voxels outside the anatomical limit of the particular regions, binary masks were created using AAL [65] for the amygdala and anterior insula, as well as resting-state network template masks for the hippocampus [66]. The time-series extracted from each ROI in the form of the first principal-variate of all the regional voxels timeseries, was used as the observed BOLD signal for the effective connectivity estimation using dynamic causal modelling.

**Table 1.**
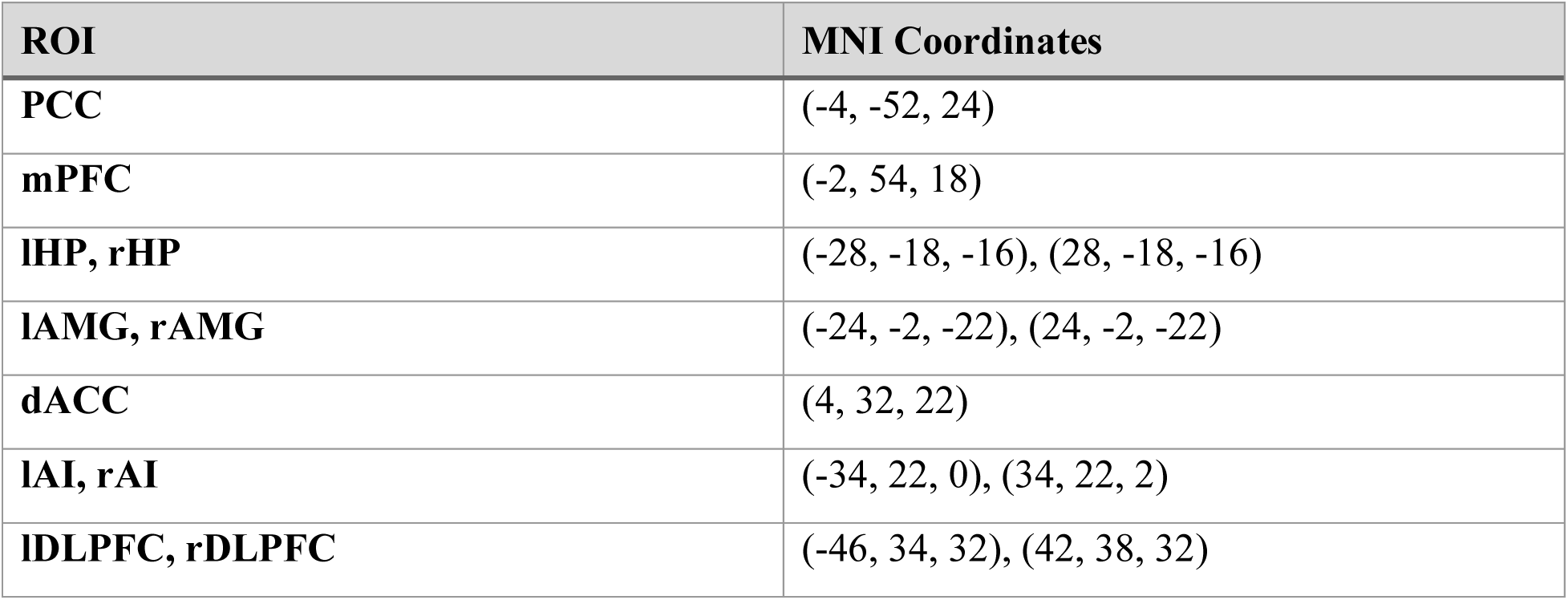
MNI Coordinates of ROIs centroids.

### Spectral dynamic causal modelling

The effective connectivity investigation for resting state fMRI of each subject was carried out using spectral DCM (spDCM; [67], [68]) - a variant of DCM to infer the resting-state or intrinsic effective connectivity among the neural population. DCM is a Bayesian framework composed of generative models of stochastic neuronal dynamics and hemodynamic response for the observed BOLD signal [69]. The fully connected spectral DCM of 11 ROIs was used to estimate the intrinsic effective connectivity using the cross-spectra of the regional timeseries. We estimated the 11 x 11 asymmetric matrix for each subject representing effective connectivity between and within networks. The average percentage of variance explained by the DCM model estimation was used to assess the accuracy of the model inversion. Please refer to the supplementary material for technical details of spDCM.

### Parametric empirical Bayes

The subjects were divided into groups (low and high) based on their self-reported emotional scores to perform association analysis of effective connectivity with each emotion and their respective neuroticism scores. The association analyses were conducted using parametric empirical Bayes (PEB; [70], [71]). Parametric empirical Bayes refers to the Bayesian hierarchical model over the parameters that explains the subject-level effects using group-level parameter estimates. The use of the entire posterior density over the parameters from each subject’s DCM is the major benefit of PEB over classical hypothesis testing which only uses the expected values of connectivity parameters, discarding the estimated parameter uncertainty. Here, the purpose of using PEB is to model the within-subject connectivity differences in association with subject-specific emotion measures. In this Bayesian GLM, a group-level variable of interest (emotional scores) and any unexplained noise defined as covariates or regressors, were used to model the subject-level parameter estimates (posterior distribution of effective connectivity). The group mean is the first regressor followed by the mean-centered emotion scores. An efficient automatic Bayesian model search was performed using Bayesian model reduction (BMR) algorithm. BMR explores the model space by pruning the connections of the full model to produce reduced models. The reduced models are examined based on log model-evidence, pruning stops when model-evidence cannot be improved further. Bayesian model averaging (BMA) is used to consider parameters from all model variations. BMA averages the models’ parameters weighted with their model evidence [72]. As a result, group-level parameters (the so-called beta-coefficients) are obtained representing the association between effective connectivity and emotional scores. For detailed information on PEB, please refer to the supplementary material.

### Leave-one-out cross-validation

The effect sizes estimated using PEB were assessed by predicting the scores using the strongest associations with leave-one-out cross-validation (LOOCV) method. This method is followed by fitting the PEB model to all the subjects except one whose scores are predicted using the estimated effects. The model was validated by quantifying the (Pearson’s) correlation between the predicted and original scores. We performed these predictions for negative emotions and neuroticism.

## Results

### Self-reported scores

The subjects were categorized into two groups—high and low—based on their emotion scores. Of the 1,079 subjects, some exhibited the lowest score in each emotional category, while the scores for the remaining subjects followed a normal distribution (Figure S1). Specifically, 40 out of 157 subjects (25.5%) had the lowest anger score of 28.6; 58 out of 103 subjects (56.3%) had the lowest fear score of 32.9; and 157 out of 237 subjects (66.2%) had the lowest sadness score of 34.2. These low self-reported scores resulted in a bimodal distribution for anger, fear, and sadness. Bouziane et al. also noted this scoring anomaly in the HCP dataset (Bouziane et al., 2022). One possible explanation for this anomaly could be the approximations used in the calculation of these scores within the HCP data analysis pipeline. However, the HCP dataset does not provide the U.S. population parameters for T-score conversion, making it impossible for us to verify and correct the scoring transformations. Following the approach of Bouziane et al., we excluded these subjects (N=176) from our analysis to mitigate the impact of potentially erroneous data (Bouziane et al., 2022).

There were 784, 786 and 773 subjects with moderate scores (60 ˃ T ˃ 40) of anger, fear and sadness respectively. Their average values were 49.48 ± 4.91, 50.78 ± 4.65, and 48.21 ± 4.58, respectively. The mean neuroticism scores of these subjects were 17.20 ± 6.75, 16.59 ± 6.57, and 17.29 ± 6.52, respectively. The moderate scores of anger, fear, and sadness were positively correlated with their respective neuroticism scores with correlation values of 0.33, 0.41, and 0.48 respectively.

There were 54 low anger scorers and 65 high anger scorers. Their mean anger-affect scores were 36.9 ± 2.38 and 64.65 ± 4.65 along with their mean neuroticism scores of 12.04 ± 5.95 and 26.74 ± 7.03 respectively. There were 17 low fear scorers and 100 high fear scorers. Their mean fear-affect scores were 37.83 ± 1.25 and 64.85 ± 4.49 along with their mean neuroticism scores of 11.53 ± 6.02 and 26.47 ± 6.55 respectively. There were 73 low sadness scorers and 57 high sadness scorers. Their mean sadness scores were 39.03 ± 0.44 and 64.69 ± 4.82 along with their mean neuroticism scores of 11.83 ± 5.87 and 28.88 ± 7.14 respectively. Please see Figure S2, S3 and S4 for data distribution visualization. Among all, there were only 19 subjects with simultaneously high scores in all three negative affect domains.

The correlation between high anger scorers and their neuroticism scores was positive with a value of 0.18 while low anger scorers were negatively correlated with their neuroticism scores with a correlation value of -0.08. High fear scorers had the strongest correlation of 0.46 with their neuroticism scores, while low fear scorers were also negatively correlated with their neuroticism scores with value -0.16. High sadness scorers also had a strong correlation of 0.35 with their neuroticism scores while low sadness scorers had a positive correlation of 0.20. Graphical illustration is shown in Figure S5.

### Individual-specific connectivity modelling

The average variance explained, or the DCM model fit to the data over all the subjects was 87.10% with minimum 50% and maximum 97% (Figure S6). These model inversion accuracies represent excellent model fitting.

### Relationship of group-specific connectivity with neuroticism

We report the results of effective connectivity association with self-reported scores of neuroticism, anger-affect, fear-affect, and sadness. Most of them were associated with self-connections. In DCM, self-connections are also called intrinsic connections or endogenous connections. These are intra-region connections while between-region connections are called extrinsic connections. In DCM, self-connection represents the rate of decay of the neural activity of the brain region and is always modelled as inhibitory. The estimated parameters of self-connections are log-scaled parameters with a default value of -0.5 Hz. They represent inhibitory influence on the region and hence control the sensitivity to the extrinsic connections or input to the region. A positive self-connection represents increased inhibition of external inputs and vice versa. The net effect of input connections from other regions decreases with the increase in the self-connection magnitude. All reported values are the estimated parameters of the averaged model that passed the threshold of posterior probability > 0.99 (representing very strong statistical evidence).

The high anger subjects have 13 effective connectivity associations with their neuroticism with self-connection of dACC as the strongest positive while the self-connection of lHP as the strongest negative one (Figure 1). The low anger subjects have 15 associations with their neuroticism scores with lAI to PCC connection as the strongest positive while lAMG to rAMG connection as the strongest negative association (Figure 1).

**Figure 1.**
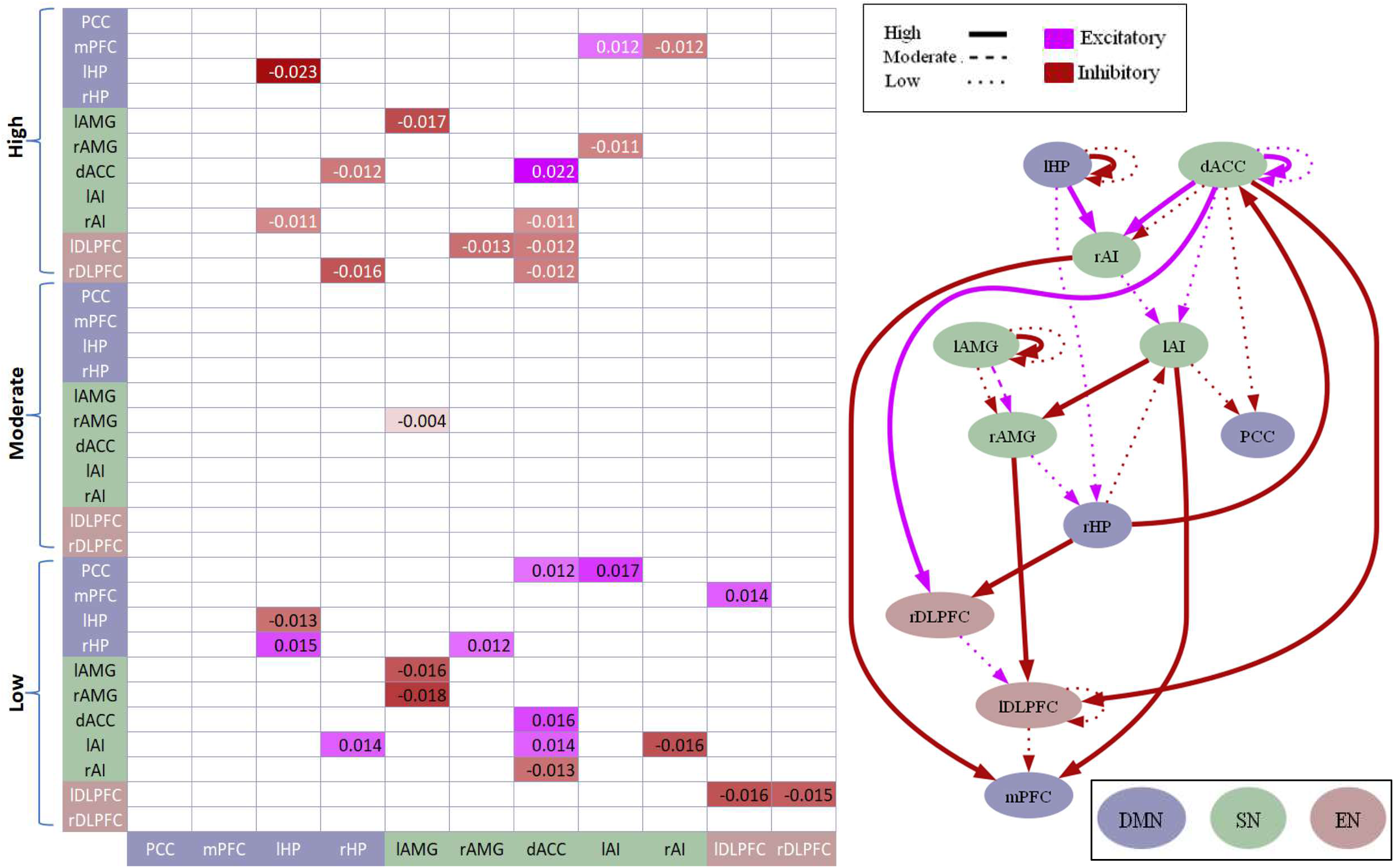
Effective connectivity association with neuroticism subjects with high, moderate and low anger. a) Matrix showing the positive and negative associations. Pink color shows positive association while blue color shows negative association. Displayed values are the normalized beta coefficients representing the contribution of group-level parameter (neuroticism) with the effective connectivity. b) The network graph showing the valence of the effective connectivity associated with the neuroticism scores of subjects with high, moderate, and low levels of anger. Different levels of emotions are represented through the line textures (thick, thin and dashed) while valence (excitation and inhibition) is represented through pink and blue colors. All values are the estimated parameters of the averaged model that passed the threshold of posterior probability > 0.99 (representing very strong statistical evidence) except moderate anger that passed the threshold of posterior probability > 0.5. The definition for label signs and threshold for estimated parameters is the same for all other figures. Abbreviations: mPFC = medial prefrontal cortex, PCC = posterior cingulate cortex, HP = hippocampus, dACC = dorsal anterior cingulate cortex, AI = anterior insula, AMG = amygdala, DLPFC = dorsolateral prefrontal cortex. Same abbreviations are used in all the figures.

The neuroticism of high fear subjects is associated with 10 effective connections. Among them the strongest positive one is the self-connection of dACC and the strongest negative is the self-connection of lAMG (Figure 2). The strongest associations of the neuroticism of low fear subjects with effective connectivity include positive association of the rHP to dACC connection and negative association of the self-connection of lHP while there are total of 31 associations (Figure 2).

**Figure 2.**
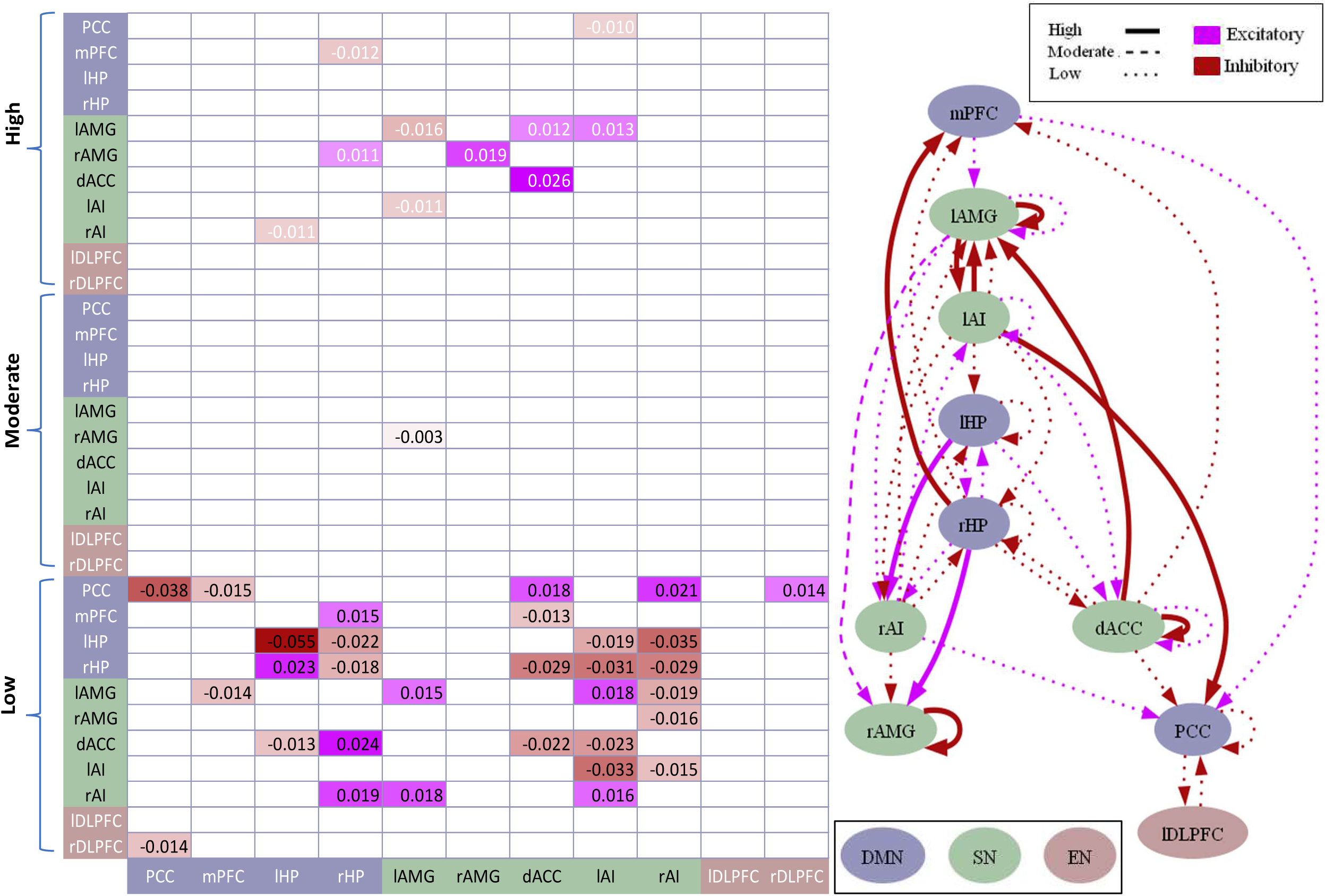
Effective connectivity association with neuroticism of subjects with high, moderate, and low fear. a) Matrix showing the positive and negative associations. Pink color shows positive association while blue color shows negative association. Displayed values are the normalized beta coefficients representing the contribution of group-level parameter (neuroticism) with the effective connectivity. b) The network graph showing the valence of the effective fear associated with the neuroticism scores of subjects with high, moderate, and low levels of anger. Different levels of emotions are represented through the line textures (thick, thin and dashed) while valence (excitation and inhibition) is represented through pink and blue colors. All values are the estimated parameters of the averaged model that passed the threshold of posterior probability > 0.99 (representing very strong statistical evidence) except moderate fear that passed the threshold of posterior probability > 0.5.

The high sadness scorers also showed associations of effective connectivity with their neuroticism scores. The strongest among 13 associations were self-connection of dACC as the positive one and the self-connection of lAMG as the negative one (Figure 3). The low sadness subjects had 13 associations. The strongest positive association was excitatory rHP to rAI the self-connection of lHP was the strongest negative association (Figure 3).

**Figure 3.**
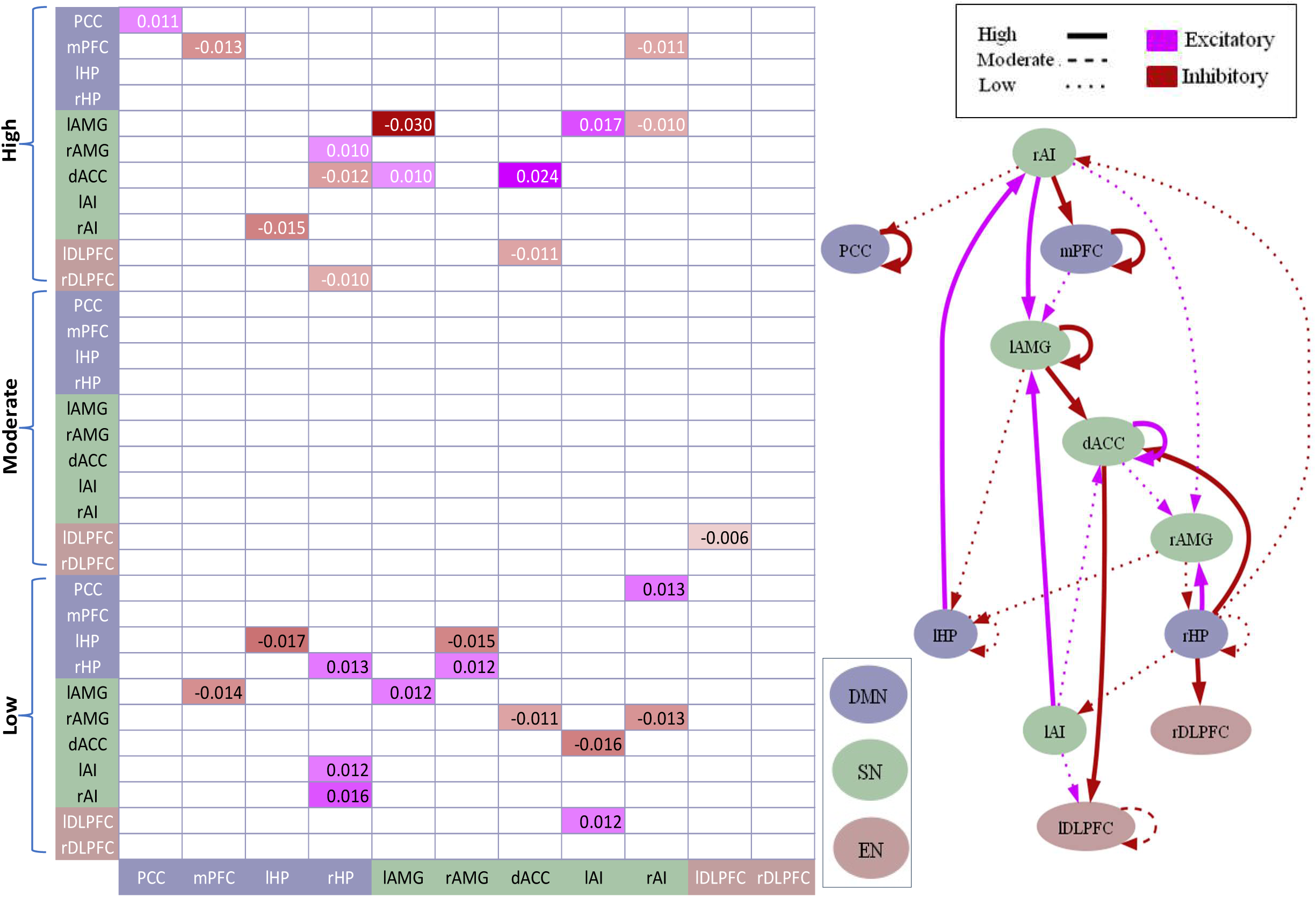
Effective connectivity association with neuroticism of subjects with high, moderate, and low sadness. a) Matrix showing the positive and negative associations. Pink color shows positive association while blue color shows negative association. Displayed values are the normalized beta coefficients representing the contribution of group-level parameter (neuroticism) with the effective connectivity. b) The network graph showing the valence of the effective connectivity associated with the neuroticism scores of subjects with high, moderate, and low levels of sadness. Different levels of emotions are represented through the line textures (thick, thin, and dashed) while valence (excitation and inhibition) is represented through pink and blue colors. All values are the estimated parameters of the averaged model that passed the threshold of posterior probability > 0.99 (representing very strong statistical evidence).

The neuroticism of moderate anger and moderate fear subjects was associated with only single connection of lAMG to rAMG negatively (Figure 1 and 2) while that of moderate sadness was associated with lDLPFC self-connection (Figure 3).

The mean connectivity of neuroticism and basic negative emotions is shown in figure S7 and S8.

### Relationship of group-specific connectivity with negative emotions

All reported results are based on the beta-coefficients that are the estimated parameters of the averaged PEB model that passed the threshold of posterior probability > 0.99 (representing very strong statistical evidence) except moderate anger and moderate fear that passed the threshold of posterior probability > 0.5. The high anger-affect was associated with 11 effective connections with strongest positive association of self-connection of lAMG and strongest negative association of inhibitory rAMG to PCC connectivity (Figure 4). The low anger-affect was also associated with 16 effective connections with strongest positive association of self-connection of rHP and strongest negative association of self-connection of rAMG (Figure 4). The high fear-affect was associated with only 4 effective connections with strongest positive association of the self-connection of rHP and strongest negative association of the self-connection of lHP (Figure 5). The low fear-affect was associated with 20 effective connections with strongest positive association of self-connection of dACC and strongest negative association of dACC to mPFC connection (Figure 5).

**Figure 4.**
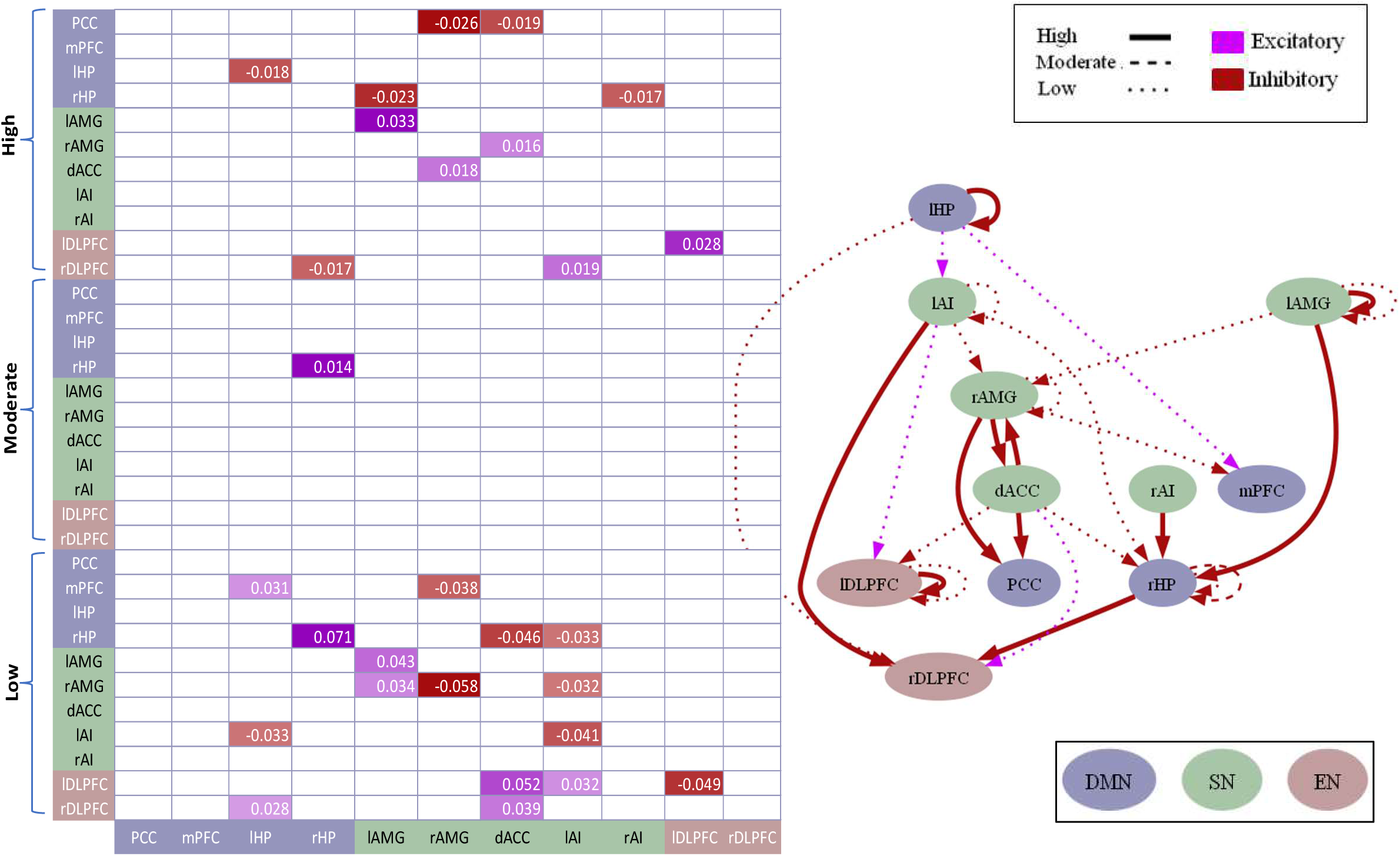
Effective connectivity association with high, moderate, and low anger. a) Matrix showing the positive and negative associations. Purple color shows positive association while red color shows negative association. Displayed values are the normalized beta coefficients representing the contribution of group-level parameter (anger) with the effective connectivity. b) The network graph showing the valence of the effective connectivity associated with high, moderate, and low levels of anger. Different levels of emotions are represented through the line textures (thick, thin, and dashed) while valence (excitation and inhibition) is represented through purple and red colors. All values are the estimated parameters of the averaged model that passed the threshold of posterior probability > 0.99 (representing very strong statistical evidence).

**Figure 5.**
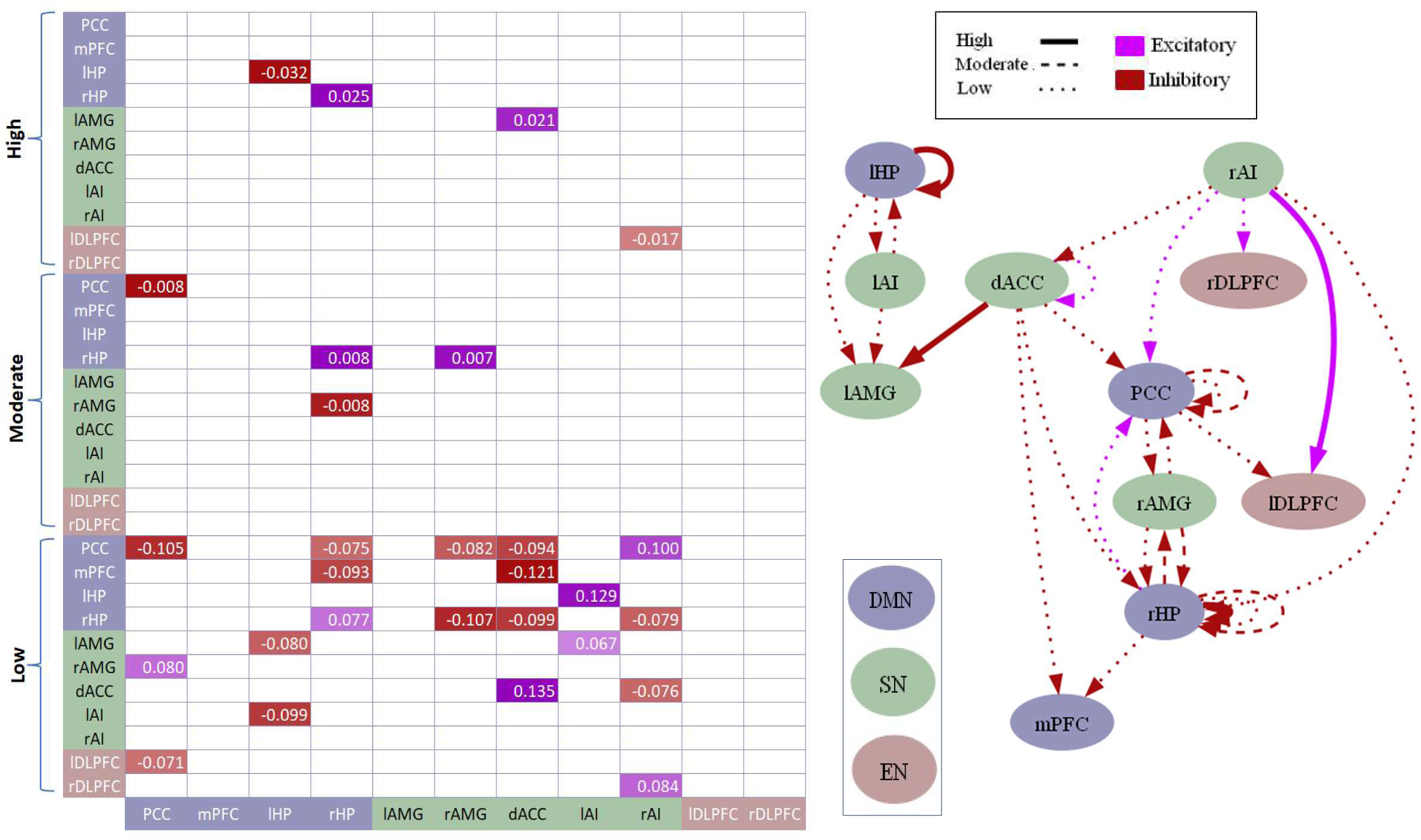
Effective connectivity association with high, moderate, and low fear. a) Matrix showing the positive and negative associations. Purple color shows positive association while red color shows negative association. Displayed values are the normalized beta coefficients representing the contribution of group-level parameter (fear) with the effective connectivity. b) The network graph showing the valence of the effective connectivity associated with high, moderate, and low levels of fear. Different levels of emotions are represented through the line textures (thick, thin, and dashed) while valence (excitation and inhibition) is represented through purple and red colors. All values are the estimated parameters of the averaged model that passed the threshold of posterior probability > 0.99 (representing very strong statistical evidence).

The high sadness was associated with 16 effective connections with strongest positive association of self-connection of dACC and strongest negative association of self-connection of lHP (Figure 6). The low sadness was associated with 9 causal connections with strongest positive association of the self-connection of lAMG and strongest negative association of lAI to rAMG connection (Figure 6).

**Figure 6.**
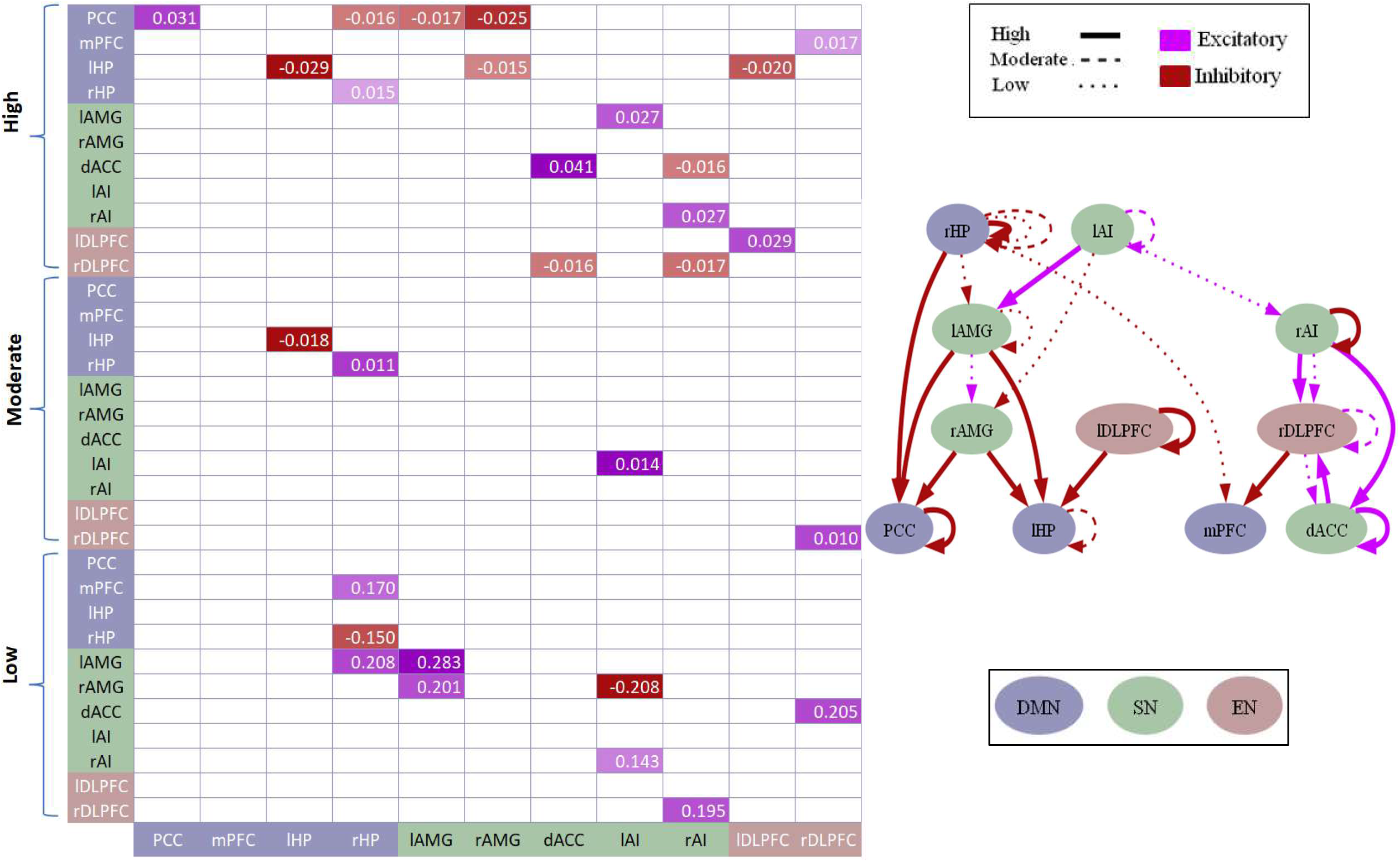
Effective connectivity association with high, moderate, and low sadness. a) Matrix showing the positive and negative associations. Purple color shows positive association while red color shows negative association. Displayed values are the normalized beta coefficients representing the contribution of group-level parameter (fear) with the effective connectivity. b) The network graph showing the valence of the effective connectivity associated with high, moderate, and low levels of sadness. Different levels of emotions are represented through the line textures (thick, thin, and dashed) while valence (excitation and inhibition) is represented through purple and red colors. All values are the estimated parameters of the averaged model that passed the threshold of posterior probability > 0.99 (representing very strong statistical evidence).

The moderate anger-affect was positively associated with the self-connection of rHP only (Figure 4). Moderate fear-affect had 4 effective connectivity associations with the strongest positive association being the self-connection of rHP while the strongest negative association being the self-connection of PCC (Figure 5). Moderate sadness was associated with 4 effective connections with self-connection of lAI as the strongest positive and self-connection of lHP as the strongest negative association (Figure 6).

### Predicting basic negative emotions and neuroticism using effective connectivity

We used the connections for prediction of self-reported scores that showed the strongest associations, using leave-one-out cross validation. We also investigated other combinations of connections for predictions such as the first two strongest positive associations or the combination of strongest positive and strongest negative associations.

The strongest positive association of high fear-affect i.e., self-connection of rHP showed significant correlation (r = 0.29, p = 0.0015) between the predicted and actual scores of high fear-affect (Figure 7(a) and Figure 8(a)). High sadness was best predicted using a combination of the top 3 positive associations (self-connections of dACC, PCC and lDLPFC) with a r = 0.27 (p = 0.022) (Figure 7(b) and Figure 8(b)). Low fear prediction gave best results with top 2 positive associations of self-connection of dACC and lAI to lHP connections (r = 0.53, p = 0.014) (Figure S9(a) and Figure S10(a)) and low anger was best predicted using combination of strongest positive and strongest negative associations of rHP and rAMG self-connections respectively (r = 0.25, p = 0.036) (Figure S9(b) and Figure S10(b)). The neuroticism score of subjects with low fear were predicted using the first two positive associations of rHP to dACC and lHP to rHP connections (r = 0.57, p = 0.008) as shown in Figure S11. Neuroticism scores of any other emotion category could not be predicted. In figure 7(a) and 7(b), all the predicted values are within the 90% credible interval (shaded region). Although the correct predictions are above chance but there is still a lot of variability to be explained. High anger and neuroticism other than low fear category could not be predicted.

**Figure 7.**
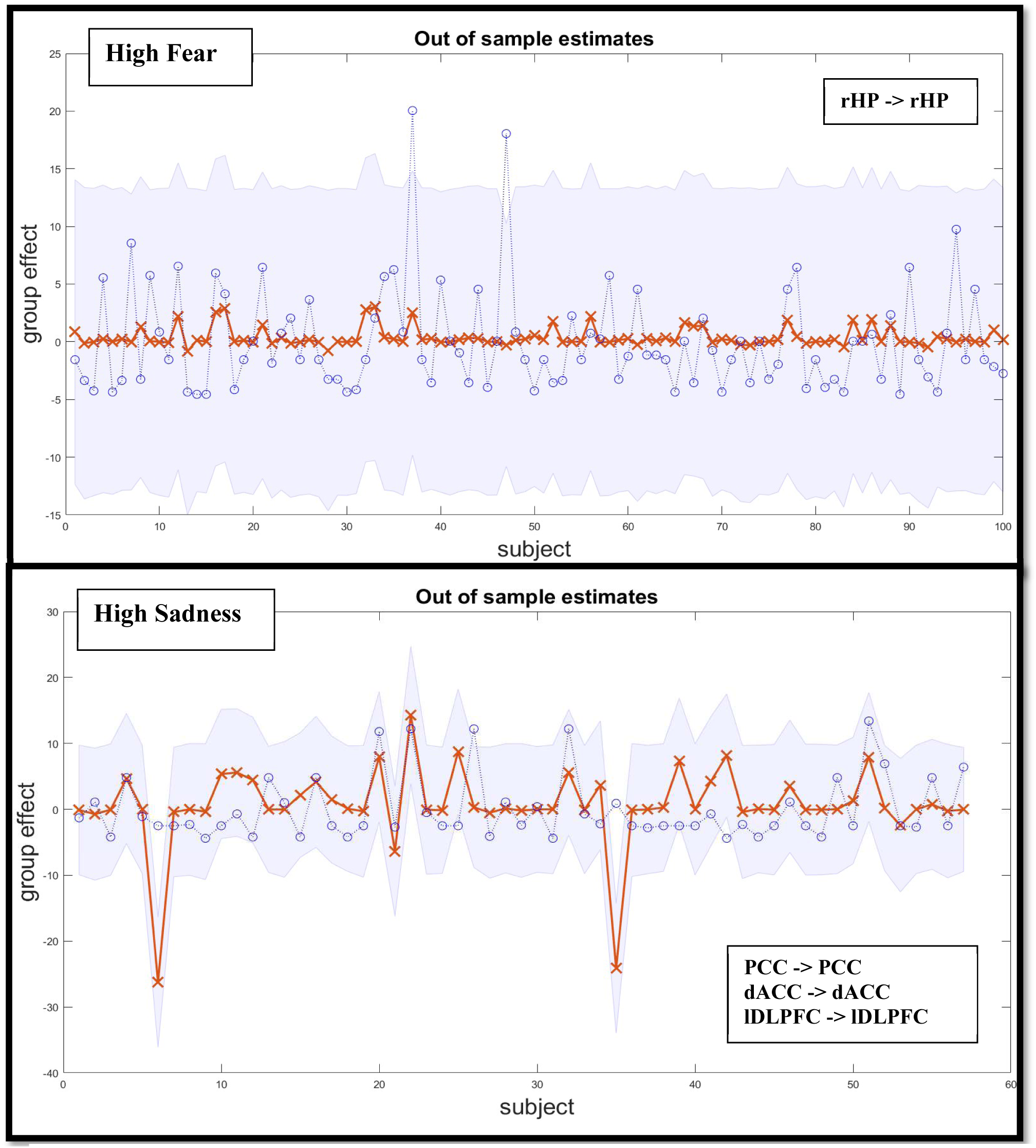
Actual and Predicted Precision in leave-one-out cross-validation of high fear and sadness. The line plot shows the predictive efficiency of the specific connections to each leave-out participant’s high fear and sadness scores. The grey area shows the 90% confidence interval of the prediction.

**Figure 8.**
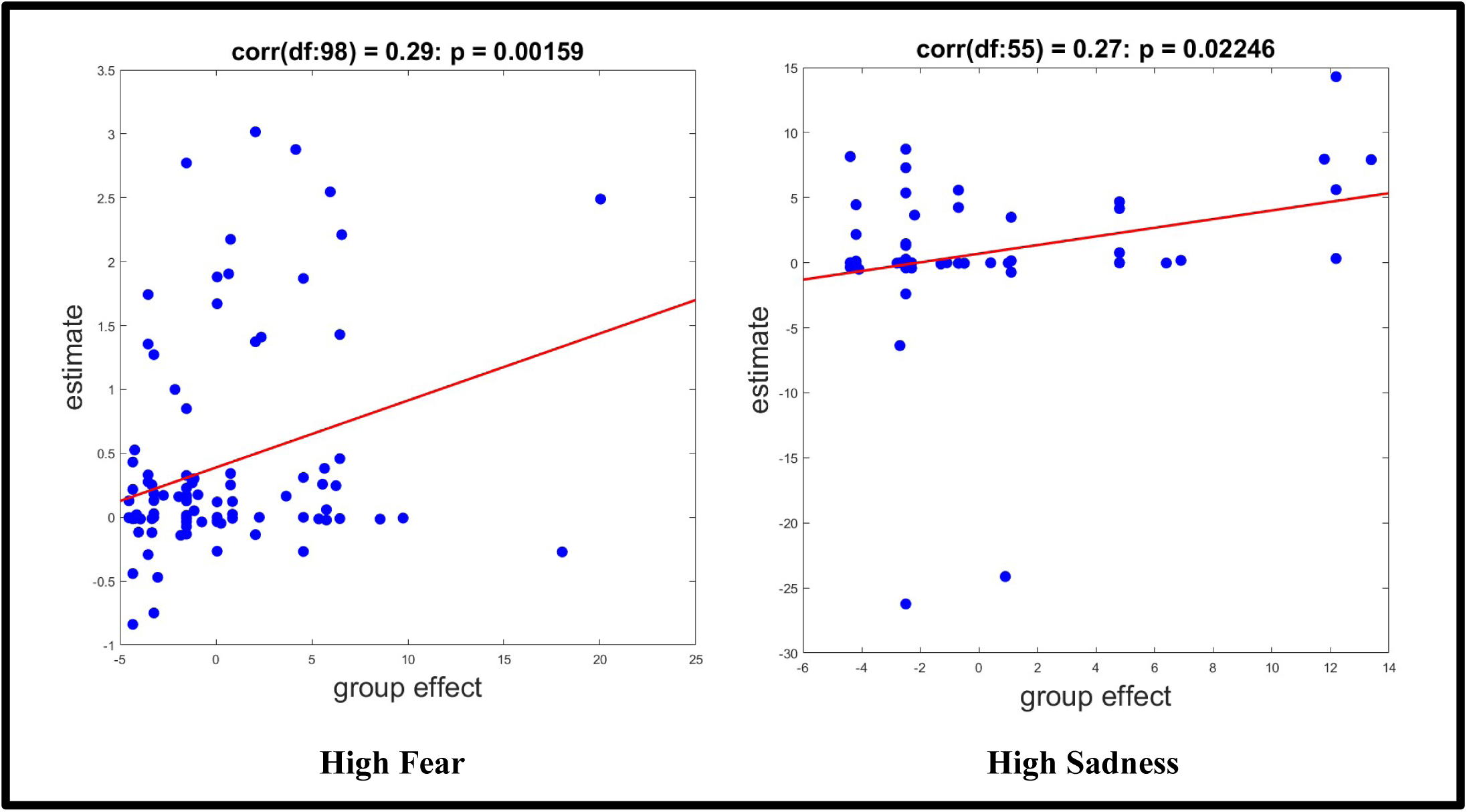
Out-of-samples correlation of high fear and low sadness. The scatterplot shows the out-of-samples Pearson’s correlation between the estimated values (estimate) and the actual values (group effect). The red line shows the linear trend fitting the dots.

The strongest positive and negative associations of effective connectivity with negative emotions and neuroticism are collectively shown in Figure 9.

**Figure 9.**
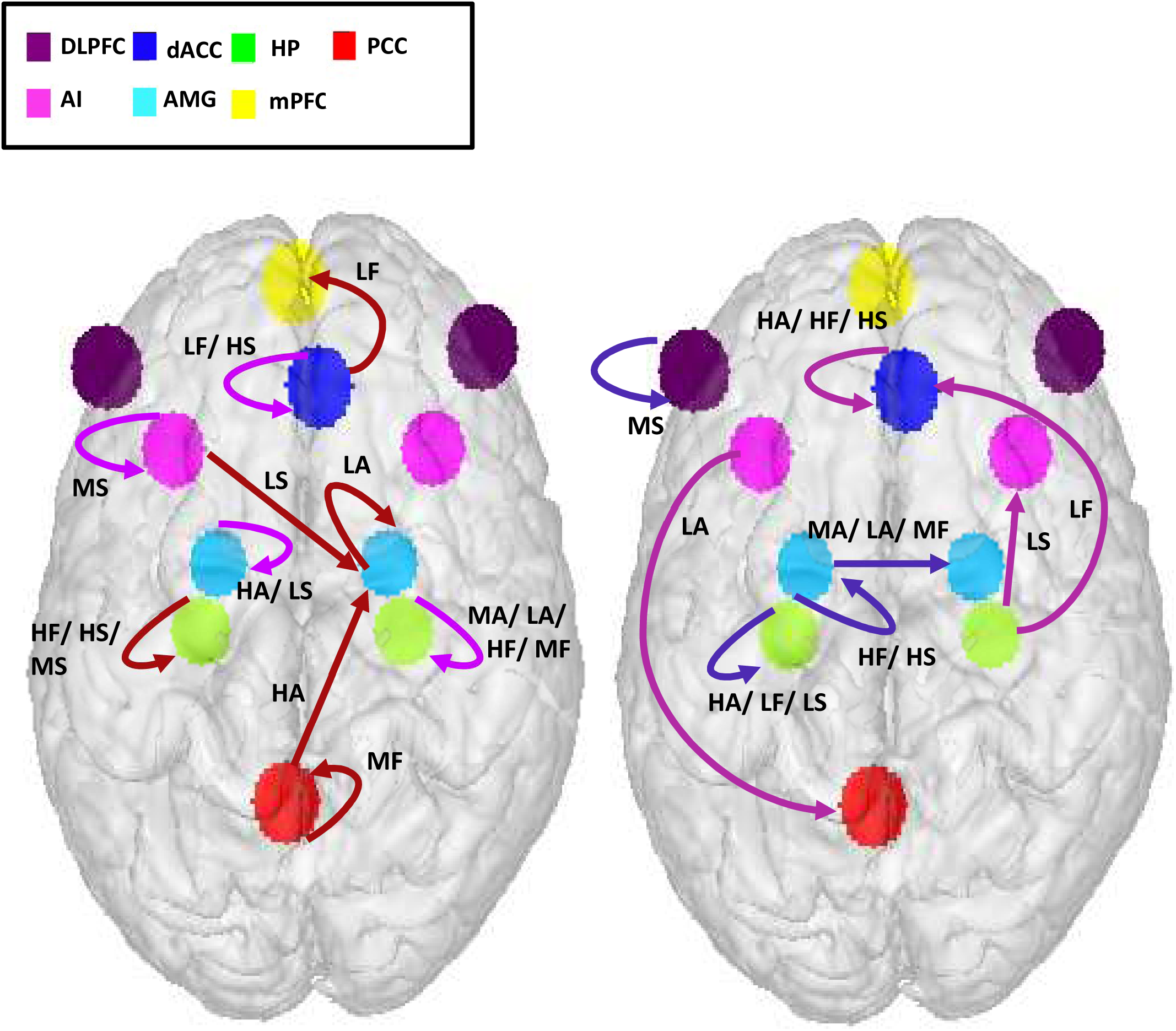
Strongest positive and strongest negative associations. a) Top-most positive and negative associations of effective connectivity with basic negative emotions. Purple and red color shows the positive and negative associations respectively. b) Top-most positive and negative associations of effective connectivity with neuroticism. Pink and blue color shows the positive and negative associations respectively. Abbreviations: HA: High anger, HF: High fear, HS: High sadness, LA: Low anger, LF: Low fear, LS: Low sadness.

## Discussion

In this study, we explored whether the effective connectivity of resting-state triple network: DMN, SN and EN, holds significance to identify neuroticism and negative affects including anger, fear, and sadness. The executive and default mode networks reciprocate each other in activity i.e., DMN is active during rest and EN during externally oriented task or vice versa [73]. The SN performs switching between these two networks. During recent past, researchers have been more focused on the interconnections of the triple network to determine if this is the seat for emotional disorders or psychopathology [74], [75]. This triple network is special because it appears to be the hub in the brain where all information including cognitive, sensory, and affective converges, interpreted, planned, and executed. We used a large-scale dataset of 1079 subjects with their self-reported scores of neuroticism and negative emotions from which 176 subjects were removed being outliers, therefore, we performed analysis on 903 subjects. Despite 85% of the subjects had moderate scores in each emotion category, sample size of low and high scores were sufficiently large (> 50) except low fear (17 subjects). While we have reported the results of low, moderate, and high levels of each emotion category but only discussed associations of high emotions scores because the previous literature is mostly focused on heightened emotions. The results revealed that the self-connections of rHP which is a part of DMN has the tendency to predict an individual with high fear while the self-connections of dACC, PCC and lDLPFC that are the regions of SN, DMN and EN respectively, are significant enough to predict high sadness. The cross-validation results are promising with above average chance of correct prediction. We found a relationship of effective connectivity with anger and neuroticism but none of those associations were strong enough in their effect sizes to significantly predict anger and neuroticism.

In a meta-analysis study, Wager et. al., revealed that a basic emotion category is not related to any single resting-state network, rather each emotion category spans across multiple cortical and subcortical networks. All basic emotions show activity across all relevant cortical and subcortical regions simultaneously such as anterior cingulate cortex, amygdala, insula, PCC, hippocampus, and prefrontal cortex, but their patterns of activations differ [76]. We also witnessed this in our association results, for example, left hippocampus was commonly associated with all high negative emotions and left amygdala and dACC was commonly related to neuroticism of all high emotions subjects.

There is already a well-established relationship between emotions and memory. The negative emotions are an outcome of self-referential and self-generated thoughts that are usually based on prior experience [77]. These thoughts are particularly significant in situations where declarative or explicit memory (episodic and/or semantic) is required to make sense of the environment. In the current study, the participant makes a conscious effort to recall memories that occurred during the last week for self-reporting of the negative emotions, hence the role of hippocampus is significant in our association results. The hippocampus has critical role in episodic memory [78]–[80] with the differential association of left and right hippocampi in verbal memory [81] and visual-spatial memory [82], [83] respectively. Previous research revealed the effect of emotions and neuropsychiatric disorders on the verbal episodic memory [84]–[86]. The verbal memory is more influenced by emotional speech, specifically negative emotions, than the neutral speech [87], [88] with enhanced activity in the left hippocampus [89]. The hippocampus is known for encoding episodic and spatial memory [90]–[92]. The critical involvement of hippocampus in spatial memory, subsequently, in the encoding and storage of contextual fear information is already established through the previous literature [93]–[95]. Fear conditioning studies revealed a distributed neural network involved in the acquisition and expression of fear, comprising amygdala, mPFC and hippocampus [95], [96]. Hippocampus has an important role in acquisition and expression of fear [97], [98]. It has been shown that the neurons reactivation in hippocampus elicits the same output as in fear conditioning, due to fear memory recall [99]. In our study, the positive association of self-connection of right hippocampus with high fear stood out as the highly promising connection to predict individuals with high fear. According to our association results, decreased inhibition of the right hippocampus will enhance high fear.

The salience network has a preferential recruitment in sadness [76] while anterior cingulate cortex and lDLPFC serves as a treatment target of sadness-related disorders such as major depression [100]. SN also recruits DMN, preferentially those regions that are involved in self-referential thoughts [76], [100], [101] specifically, mPFC and PCC were found to be related to major depression in a study [102]. The posterior regions of DMN including PCC are particularly related to the episodic memory retrieval [103], [104]. The PCC is well-acknowledged for its contribution in social cognition [105], self-reflection [106] and interaction mechanisms between memory and emotions [107]. Among the DMN regions, self-referential processes are driven by PCC activity while modulated by mPFC [55]. The negative affect is typically related to exaggerated self-referential activity, the overindulgence of one’s thoughts about the self may lead to the negative interpretation of the situations. Our results indicate decreased inhibition of PCC or increased influence on PCC from other brain regions to be positively associated with high sadness. The dACC, being the key node of the salience network integrates information from the internal and external sources to guide the behavior. The dACC has been important in several chronic sadness related studies such as depression [108], [109]. It is also considered as a good predictor of treatment response in depression [110]–[112]. Our results imply increased inhibition of dACC or decreased influence on dACC from other brain regions to be positively associated with high sadness. The DLPFC, as a component of EN, is engaged during cognitively demanding tasks [113]. One of the functions of DLPFC is to perform top-down regulation for appropriate behavioral response [114], [115] and cognitive regulation of negative emotions [116]–[121]. Hypoactivation of lDLPFC was found to be linked with negative emotional judgment in major depression [122], [123]. It has also been effectively used as stimulation target for non-invasive treatment techniques for major depression [124], [125]. Our results indicate decreased inhibition of lDLPFC or increased influence on lDLPFC from other brain regions to be positively associated with high sadness. The findings of the current study emphasized the importance of PCC, dACC and lDLPFC in the connectome-based identification of individuals with high sadness.

Among the negative emotions, only anger could not be predicted using cross-validation. The reasons could be the mutual interaction of the other two negative emotions with anger. In our dataset, there were a total of 65 high anger scorers. Out of those, 30 (almost half) had mutually high fear and high anger, 27 (almost half) had mutually high sadness and high anger while 19 had high levels of all three negative emotions. According to the previous research, anger is strengthened by fear and weakened by sadness [126], [127]. This might be the case that our anger sample was influenced by other two emotions, hence, could not be predicted through the effective connectivity associations as they have a share of other two feelings. In a meta-analysis of 148 studies, Wager et. al., also tried to predict basic emotions using functional brain patterns, their prediction results also varied with fear bearing the highest prediction accuracy while anger prediction being the lowest accurate [76].

In our study, neuroticism associations with effective connectivity revealed the strongest positive association of dACC self-connection in individuals with high levels of all three negative emotions. Self-reported neuroticism is correlated with dACC activity and neural correlates of dACC are a good predictor of neuroticism than self-reports according to the previous research [128]. There is a vast literature available providing evidence for the role of amygdala in neuroticism. The increased resting-state effective connectivity from right amygdala to right middle frontal gyrus and decreased effective connectivity from right precuneus to right amygdala was found to be associated with neuroticism [23], right amgydala-prefrontal and left amygdala-dACC functional connectivity was found to be positively and negatively associated with neuroticism in a gender identification of scrambled pictures viewing task [129], increase in resting-state functional connectivity of amygdala-precuneus and decrease in amygdala with temporal regions and insula was associated with high neuroticism [130], impaired functional connectivity between amygdala and frontal networks was also found in highly neurotic subjects during emotion regulation while avoiding negative pictures [26], individuals with higher risk of genetically-transferred neuroticism showed decreased activity in left amygdala [131]. In our results, the negative associations of the self-connection of left amygdala with neuroticism was the most prominent in individuals with high levels of fear and sadness. It suggests that decreasing inhibition of left amygdala or increasing sensitivity to other regions’ inputs might decrease neuroticism in highly feared and highly sad individuals.

We found association of triple network effective connectivity with neuroticism but the effect sizes of the strongest associations failed to identify left out individual neuroticism score in cross-validation. The reason could be that the causal interaction of triple network has not much influence on the neurotic personality type. It is also stated previously that the frequency of negative emotions is the strong predictor of neuroticism in comparison to their duration [33] therefore, the aspects of frequency and duration of emotions should also be taken into consideration for neuroticism prediction. In our dataset, the neuroticism was correlated with high anger with correlation coefficient of r = 0.18, with high sadness as r = 0.35 and with high fear as r = 0.46. The individuals with high fear and high sadness were strongly correlated with neuroticism than those with high anger.

## Conclusion

Our key findings revealed that the effective connectivity of large-scale resting-state triple network can predict high sadness and fear, specifically the self-connection of right hippocampus and self-connections of dorsal anterior cingulate cortex, posterior cingulate cortex and dorsolateral prefrontal cortex can be the biomarkers or signature brain connections for individuals with high fear and high sadness respectively. The causal interconnections of DMN, SN and EN are not good predictors of high anger, low sadness, and neuroticism of any emotion category except low fear.

Gender and age have an impact on brain connectivity and there is some bias in emotional processing and personality which was not specifically controlled for this study. Previously, the HCP data was used to predict the personality based on functional connectivity of nine resting-state networks and the results implies that personality predictions are gender-specific, most networks predicted personality in either of male or female group [46]. Future studies should explore the gender-specific predictions of neuroticism and basic negative emotions using effective brain connectivity. The influence of negative emotions on each other and specifically influence of fear and sadness on anger, should also be considered for the prediction analysis.

## Acknowledgment

Data were provided [in part] by the Human Connectome Project, WU-Minn Consortium (Principal Investigators: David Van Essen and Kamil Ugurbil; 1U54MH091657) funded by the 16 NIH Institutes and Centers that support the NIH Blueprint for Neuroscience Research; and by the McDonnell Center for Systems Neuroscience at Washington University.

