## Supplementary Information for "Neuroticism and emotion regulation: An effective connectivity analysis of large-scale resting-state brain networks"

### Dynamic causal modelling

The Effective connectivity investigation for resting state fMRI of each subject was carried out using spectral DCM ((spDCM; Friston et al., 2014) - a variant of DCM to infer the resting-state or intrinsic effective connectivity among the neural population in the absence of external stimuli. DCM is a Bayesian framework composed of generative models of stochastic neuronal dynamics and hemodynamic response for the observed BOLD signal. These generative models are inverted using Bayesian model inversion to estimate the neuronal and hemodynamic parameters that expose the hidden neuronal state causing the BOLD signal. The two generative models can be represented mathematically as,

$$\dot{x}(t) = f(x(t), u(t), \theta) + v(t), \tag{S1}$$

$$y(t) = h(x(t), \varphi) + e(t), \tag{S2}$$

where x represents the regional latent neuronal state, $\dot{x}(t)$ defines the differential equation representing change in the state of neuronal state with respect to time, u represents the external input which would be zero for resting state. A non-linear hemodynamic response function h maps the neuronal state x to the observed BOLD signal. The parameters $\theta$ and $\varphi$ (based on the Balloon model (Stephan et al., 2007) are the estimates for the neuronal state and hemodynamic response functions respectively while v and e represents the endogenous or intrinsic neural fluctuations and observation noise respectively.

The spectral DCM is intended to estimate the intrinsic effective connectivity from resting state fMRI using the cross-spectra of the signals. The cross spectral in the frequency domain, being the Fourier transform of the cross-correlation, can be considered as a measure of functional connectivity (Park et al., 2018) The endogenous fluctuations and observation noise is modelled using power law (scale free form) (Beggs & Plenz, 2003; Shin & Kim, 2006; Stam & De Bruin, 2004) as follows:

$$g_v(\omega, \theta) = \alpha_v \omega^{-\beta v} \quad \text{(S3)}$$

$$g_e(\omega, \theta) = \alpha_e \omega^{-\beta e}$$

The full model is then inverted using Variational Laplace. We estimated the 11 x 11 non-symmetric matrix for each subject representing effective connectivity between and within regions using spectral DCM.

**Parametric empirical Bayes**

Parametric empirical Bayes for DCM refers to the Bayesian hierarchical model over the parameters that explains the subject-level effects using group-level parameter estimates (Karl J. Friston et al., 2016; Zeidman et al., 2019). It is a better approach for group-level analysis in comparison to classical methods like ANOVA because it takes the complete posterior density into consideration, hence incorporating the variance within and between subjects.

Empirical Bayes refers to the Bayesian inversion or fitting of hierarchical models. In hierarchical models, constraints on the posterior density over model parameters at any given level are provided by the level above. These constraints are called empirical priors because they are informed by empirical data.

PEB for DCM is based on Bayesian GLM in which subject-level connectivity parameters are modelled using group-level parameters and some random noise. It can be used to test hypothesis on group parameters by comparing the evidence of different models using Bayesian model comparison. PEB uses the Bayesian Model Reduction (BMR) algorithm to invert many models from a single dataset or a single (hierarchical) model from numerous datasets. Mathematically, for DCM studies with N subjects and M parameters per DCM, we have a hierarchical model, where the responses of the i-th subject and the distribution of the parameters over subjects can be modeled as:

$$y_i = \Gamma_i^{(1)}(\theta^{(1)}) + \varepsilon_i^{(1)} \quad \text{(S4)}$$

$$\theta^{(1)} = \Gamma^{(2)}\left(\theta^{(2)}\right) + \varepsilon^{(2)}$$

$$\theta^{(2)} = \eta + \varepsilon^{(3)}$$

where, $y_i$ is the BOLD time series from i-th subject and $\Gamma_i^{(1)}$ is a nonlinear mapping from the parameters of a model to the predicted response $y$ for e.g. as shown in Eq. S1 above. $\varepsilon_i^{(1)}$ is independent and identically distributed (i.i.d.) observation noise (equivalent to $e(t)$ in Eq. S2). In this hierarchical form, empirical priors encoding second (between-subject) level effects place constraints on subject-specific parameters.

The between-subject part encodes differences among subjects or covariates such as age, while the within-subject part specifies mixtures of parameters that show random effects. We assume that the first column of the design matrix is a constant term, modelling group means and subsequent columns encode group differences or covariates such as age.

### Figure legend

**Figure S1.** Plot of probability density function of the scores of anger-affect, fear-affect and sadness of 1079 subjects.

**Figure S2.** Graph showing data sample size with male and female count represented through blue and pink bars respectively.

**Figure S3.** Correlation of negative emotions with their neuroticism scores.

**Figure S4.** Plot of probability density function of the scores of anger-affect, fear-affect and sadness of 903 subjects.

**Figure S5.** Plot of probability density function of the scores of neuroticism scores of 903 subjects with included in each emotion category.

**Figure S6.** Accuracy of DCM model estimation.

**Figure S7. Mean effective connectivity matrices of self-reported negative emotions.** Mean effective connectivity matrix of self-reported (a) high anger-affect (b) low anger-affect (c) high fear-affect (d) low fear-affect (e) high sadness (f) low sadness. In all figures, positive gradient represents excitatory connections while negative represents inhibitory connections except the diagonal that represents the self-connections (inhibitory by default). All values are effect sizes (posterior expectations) of connections in units of Hz except self-connections which are always modeled inhibitory and are log-scaled.

**Figure S8. Mean effective connectivity matrices of neuroticism of each basic negative emotion category.** Mean effective connectivity matrix of self-reported neuroticism scores of subject group with (a) high anger-affect (b) low anger-affect (c) high fear-affect (d) low fear-affect (e) high sadness (f) low sadness. In all figures, positive gradient represents excitatory connections while negative represents inhibitory connections except the diagonal that represents the self-connections (inhibitory by default). All values are effect sizes (posterior expectations) of connections in units of Hz except self-connections which are always modeled inhibitory and are log-scaled.

**Figure S9. Actual and predicted precision in leave-one-out cross-validation of low anger and fear.** The line plot shows the predictive efficiency of the specific connections to each leave-out participant's low anger and fear scores. The grey area shows the 90% confidence interval of the prediction.

**Figure S10. Out-of-samples correlation of low anger and fear.** The scatterplot shows the out-of-samples Pearson's correlation between the estimated values (estimate) and the actual values (group effect) of low anger and fear. The red line shows the linear trend fitting the dots.

**Figure S11. Neuroticism prediction of subjects with low fear.** The line plot shows the predictive efficiency of the specific connections to each leave-out participant's neuroticism who had low fear. The grey area shows the 90% confidence interval of the prediction. The scatterplot shows the out-of-samples Pearson's correlation between the estimated values (estimate) and the actual values (group effect) of neuroticism of subjects with low fear. The red line shows the linear trend fitting the dots.

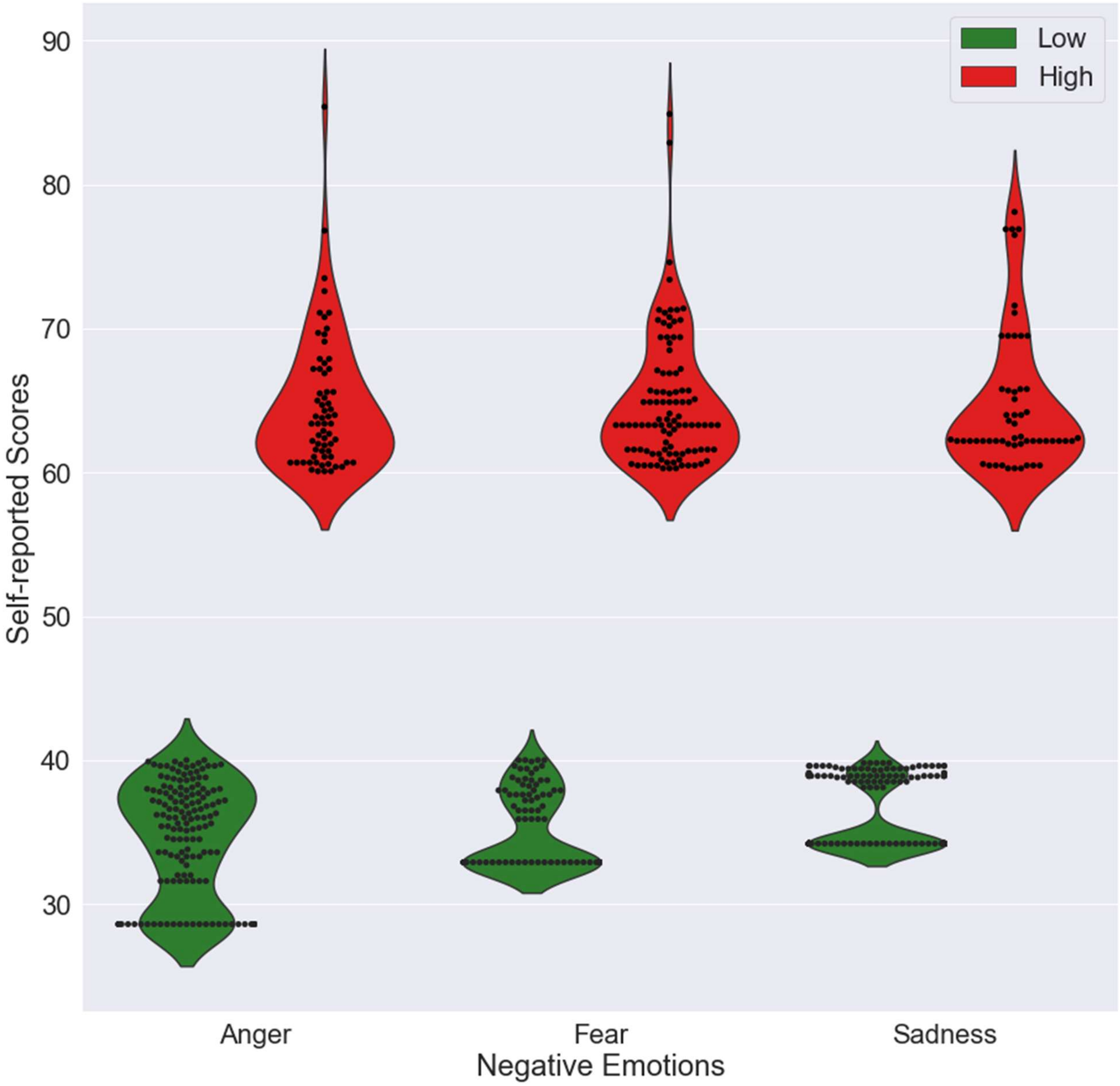

Low
High
90
80
70
60
50
40
30
Self-reported Scores
Anger
Fear
Sadness
Negative Emotions


**Figure S1**

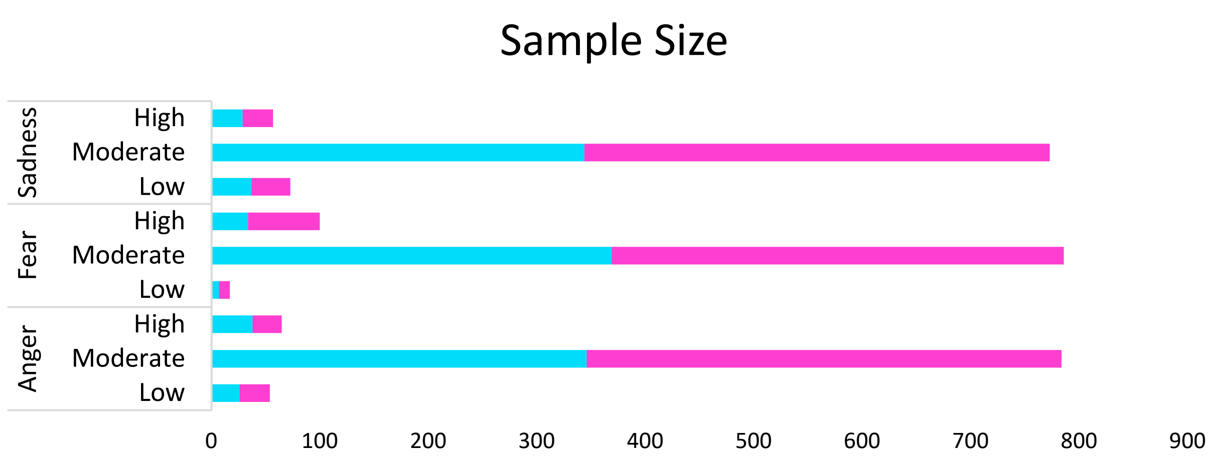


| | Anger | | | Fear | | | Sadness | | |
|---|---|---|---|---|---|---|---|---|---|
| | Low | Moderate | High | Low | Moderate | High | Low | Moderate | High |
| Male | 26 | 346 | 38 | 7 | 369 | 34 | 37 | 344 | 29 |
| Female | 28 | 438 | 27 | 10 | 417 | 66 | 36 | 429 | 28 |

**Figure S2**

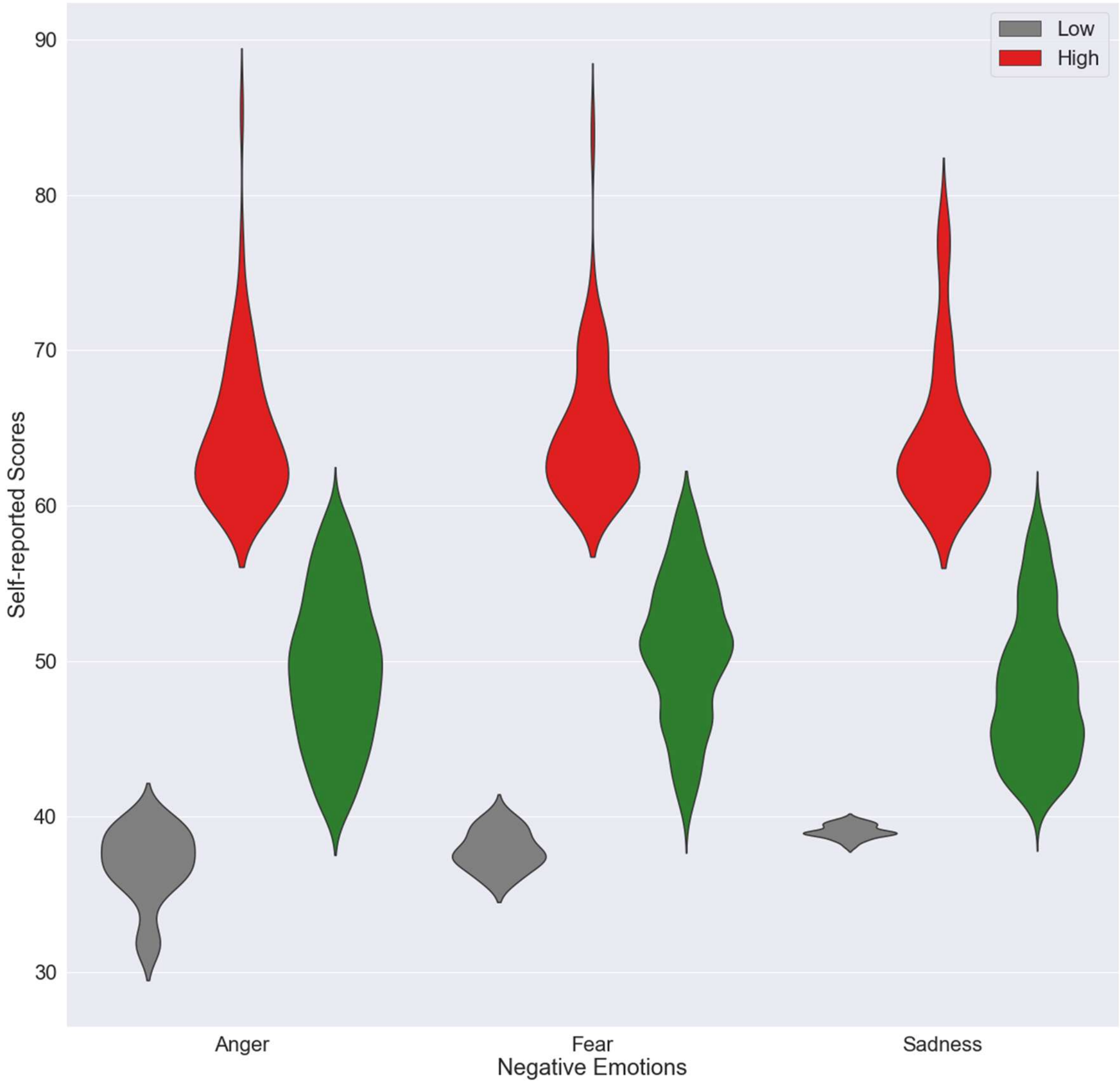
Low
High
Self-reported Scores
90
80
70
60
50
40
30
Anger
Fear
Sadness
Negative Emotions

**Figure S3**

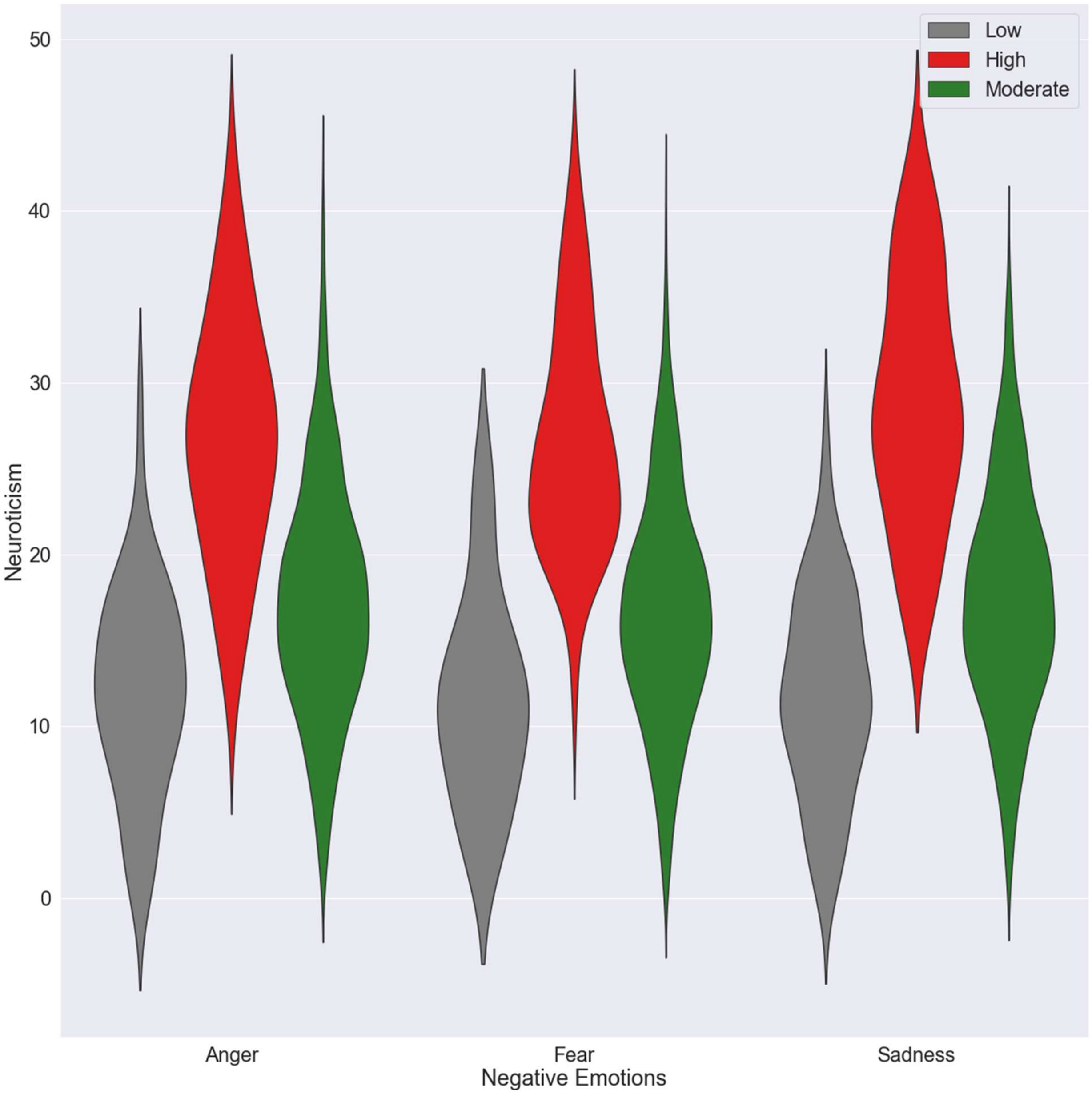
Low
High
Moderate
50
40
30
20
10
0
Neuroticism
Anger
Fear
Sadness
Negative Emotions
**Figure S4**

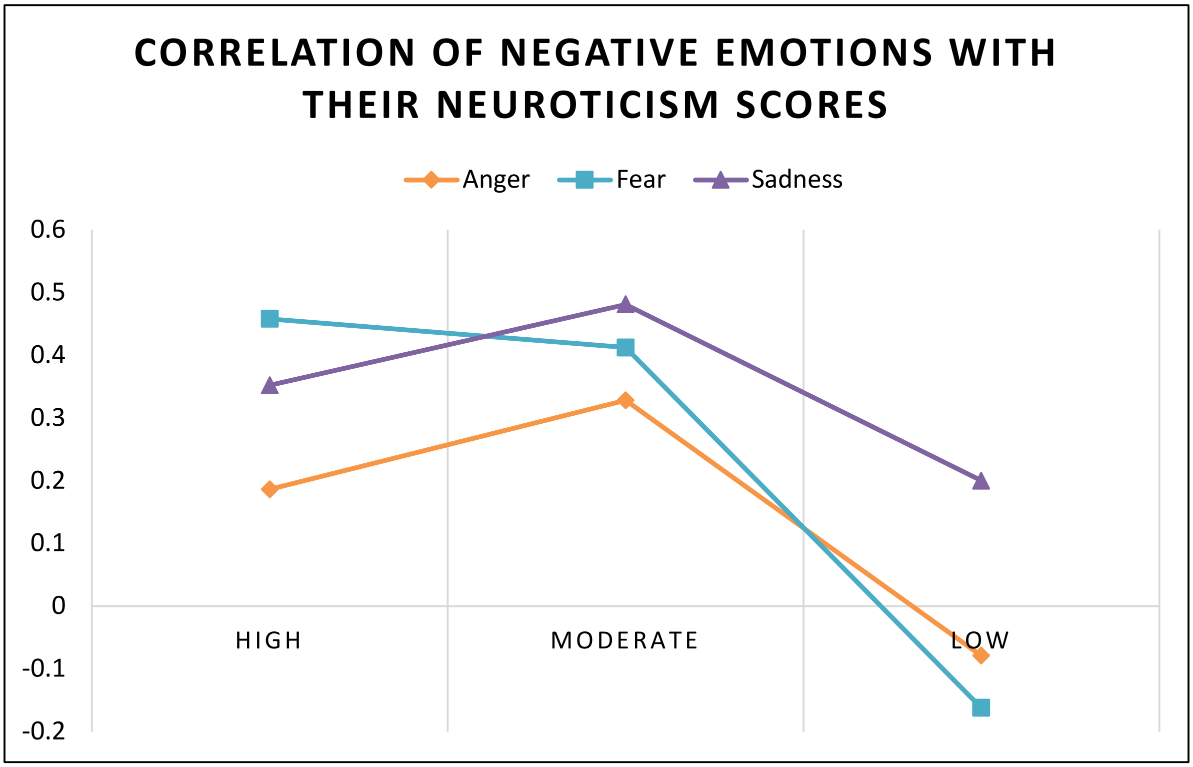
CORRELATION OF NEGATIVE EMOTIONS WITH THEIR NEUROTICISM SCORES
Anger
Fear
Sadness
0.6
0.5
0.4
0.3
0.2
0.1
0
-0.1
-0.2
HIGH
MODERATE
LOW

**Figure S5**

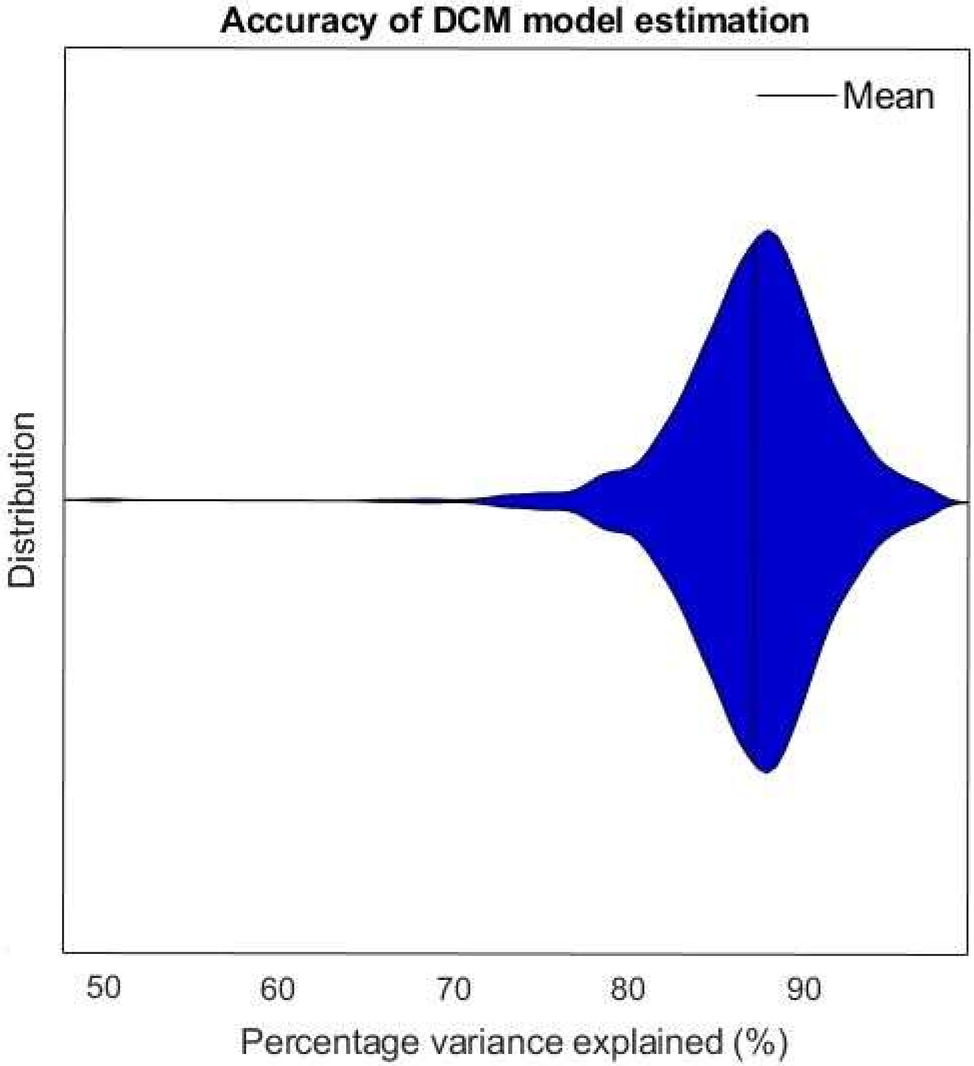
Accuracy of DCM model estimation
Mean
Distribution
50
60
70
80
90
Percentage variance explained (%)

**Figure S6**

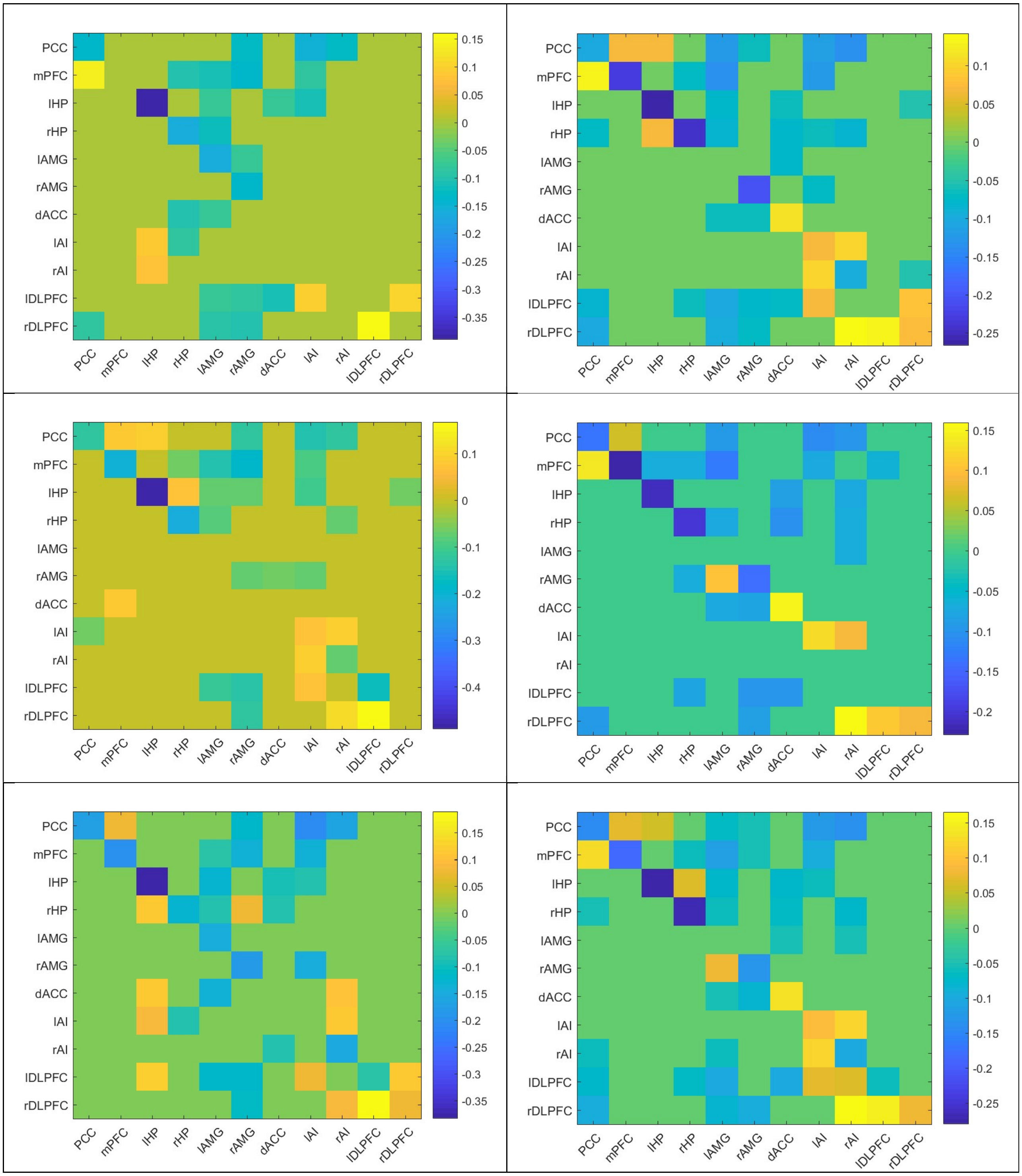


**Figure S7**

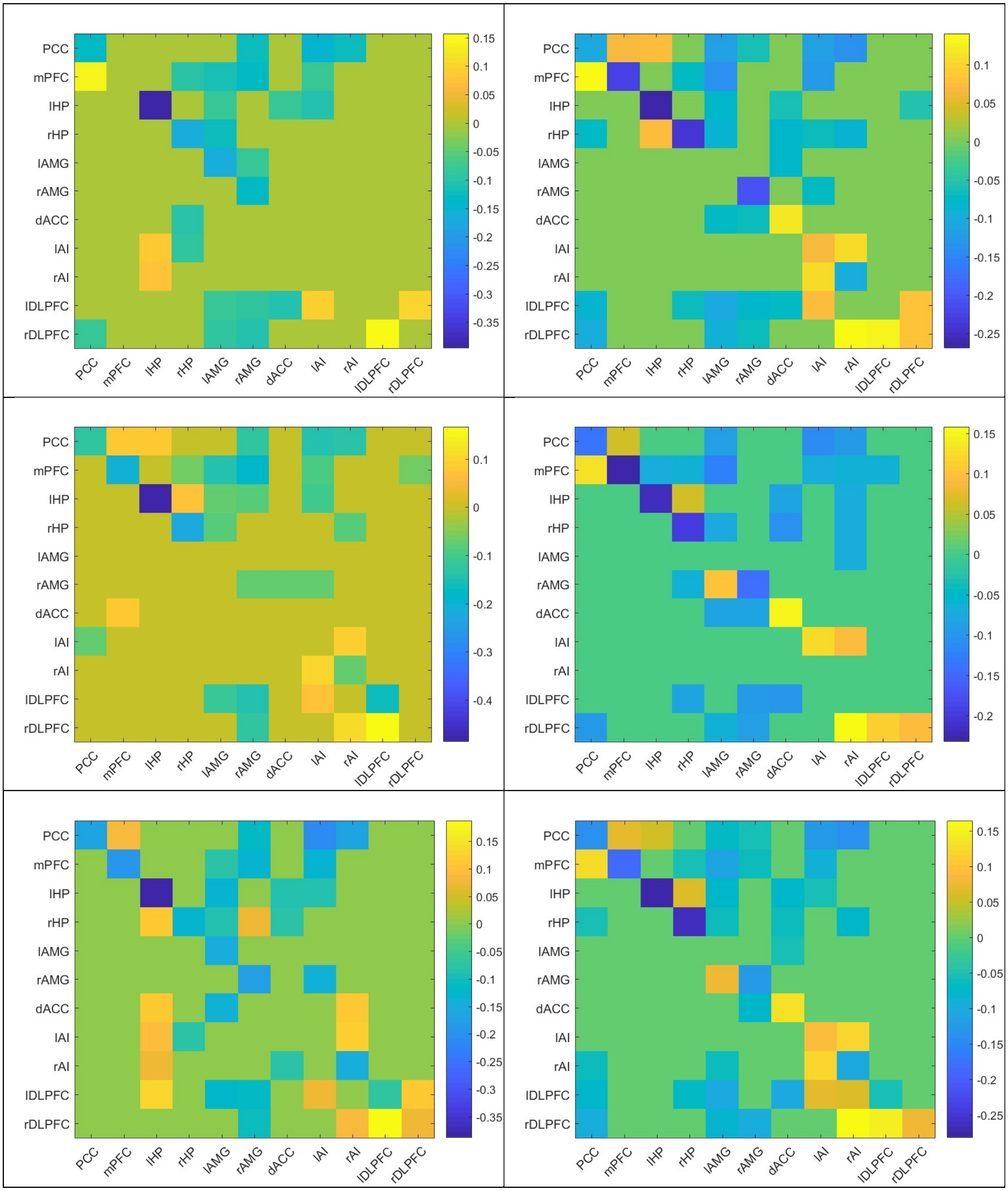
PCC
mPFC
lHP
rHP
lAMG
rAMG
dACC
lAI
rAI
lDLPFC
rDLPFC
0.15
0.1
0.05
0
-0.05
-0.1
-0.15
-0.2
-0.25
-0.3
-0.35
PCC
mPFC
lHP
rHP
lAMG
rAMG
dACC
lAI
rAI
lDLPFC
rDLPFC
0.1
0.05
0
-0.05
-0.1
-0.15
-0.2
-0.25
PCC
mPFC
lHP
rHP
lAMG
rAMG
dACC
lAI
rAI
lDLPFC
rDLPFC
0.1
0
-0.1
-0.2
-0.3
-0.4
PCC
mPFC
lHP
rHP
lAMG
rAMG
dACC
lAI
rAI
lDLPFC
rDLPFC
0.15
0.1
0.05
0
-0.05
-0.1
-0.15
-0.2
PCC
mPFC
lHP
rHP
lAMG
rAMG
dACC
lAI
rAI
lDLPFC
rDLPFC
0.15
0.1
0.05
0
-0.05
-0.1
-0.15
-0.2
-0.25
-0.3
-0.35
PCC
mPFC
lHP
rHP
lAMG
rAMG
dACC
lAI
rAI
lDLPFC
rDLPFC
0.15
0.1
0.05
0
-0.05
-0.1
-0.15
-0.2
-0.25

**Figure S8**

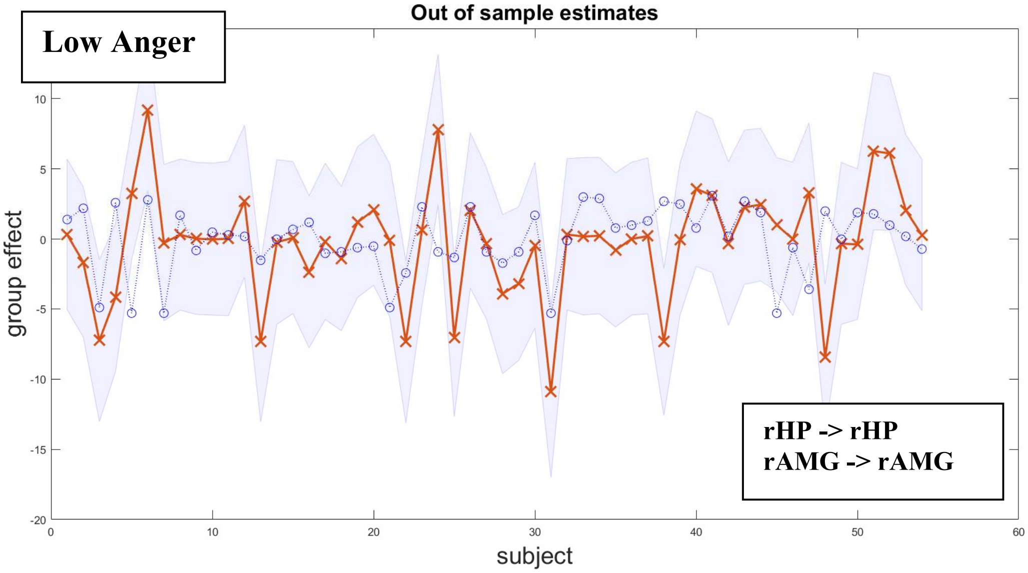
Low Anger
Out of sample estimates
rHP -> rHP
rAMG -> rAMG
group effect
subject
10
5
0
-5
-10
-15
-20
0
10
20
30
40
50
60

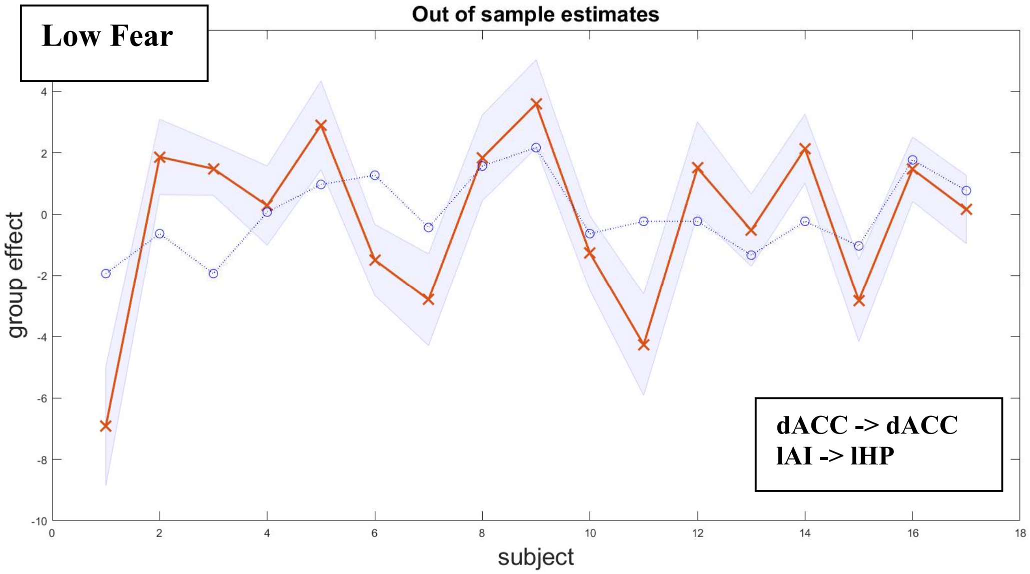
Low Fear
Out of sample estimates
dACC -> dACC
lAI -> lHP
group effect
subject
4
2
0
-2
-4
-6
-8
-10
0
2
4
6
8
10
12
14
16
18

**Figure S9**

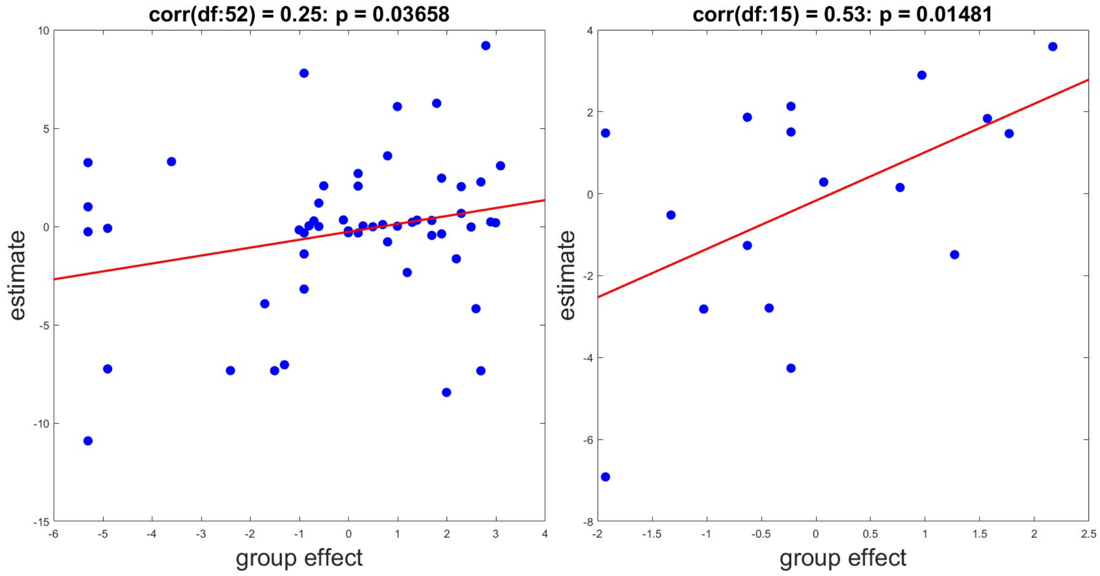
corr(df:52) = 0.25: p = 0.03658
estimate
group effect
corr(df:15) = 0.53: p = 0.01481
estimate
group effect

**Figure S10**

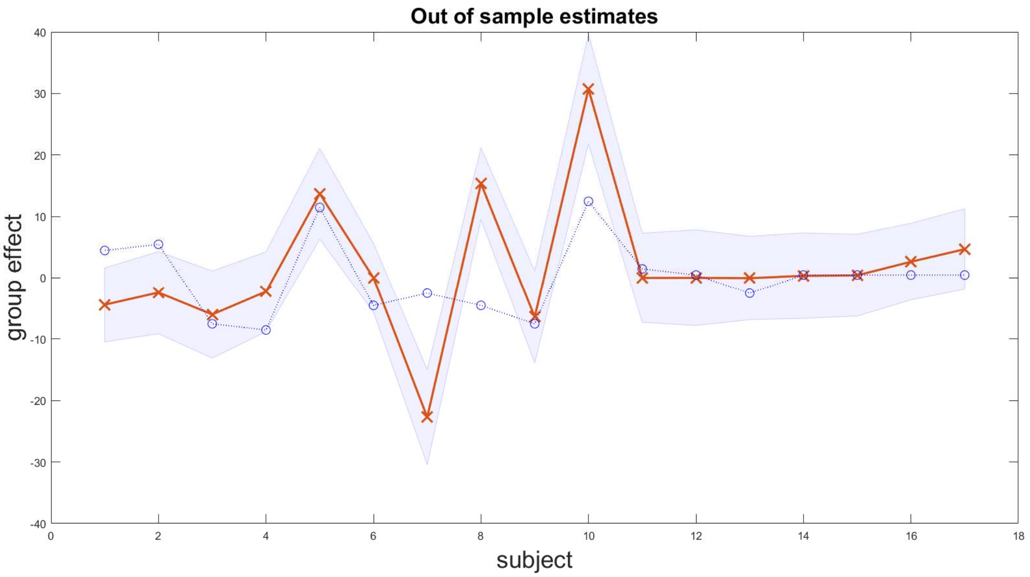
Out of sample estimates
group effect
subject
40
30
20
10
0
-10
-20
-30
-40
0
2
4
6
8
10
12
14
16
18

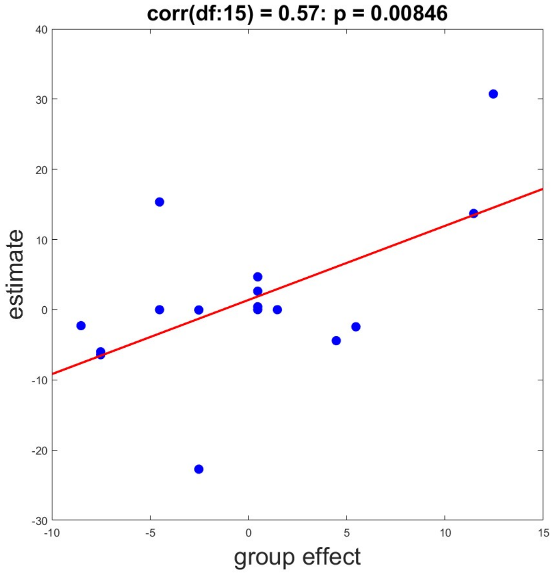
corr(df:15) = 0.57: p = 0.00846
estimate
group effect
40
30
20
10
0
-10
-20
-30
-10
-5
0
5
10
15

**Figure S11**
